# Quantifying multi-layered expression regulation in response to stress of the endoplasmic reticulum

**DOI:** 10.1101/308379

**Authors:** Justin Rendleman, Zhe Cheng, Shuvadeep Maity, Nicolai Kastelic, Mathias Munschauer, Kristina Allgoewer, Guoshou Teo, Yun Bin Zhang, Amy Lei, Brian Parker, Markus Landthaler, Lindsay Freeberg, Scott Kuersten, Hyungwon Choi, Christine Vogel

## Abstract

The mammalian response to endoplasmic reticulum (ER) stress dynamically affects all layers of gene expression regulation. We quantified transcript and protein abundance along with footprints of ribosomes and non-ribosomal proteins for thousands of genes in cervical cancer cells responding to treatment with tunicamycin or hydrogen peroxide over an eight hour time course. We identify shared and stress-specific significant regulatory events at the transcriptional and post-transcriptional level and at different phases of the experiment. ER stress regulators increase transcription and translation at different times supporting an adaptive response. ER stress also induces translation of genes from serine biosynthesis and one-carbon metabolism indicating a shift in energy production. Discordant regulation of DNA repair genes suggests transcriptional priming in which delayed translation fine-tunes the early change in the transcriptome. Finally, case studies on stress-dependent alternative splicing and protein-mRNA binding demonstrate the ability of this resource to generate hypotheses for new regulatory mechanisms.

## Introduction

Each mammalian cell coordinates the synthesis and degradation of millions of RNA and protein molecules every single minute (Harper et al., 2016) – and the relationship between transcription, translation, and degradation is highly complex (Liu et al., 2016; Schwanhäusser et al., 2011; Vogel and Marcotte, 2012). A single gene locus in DNA can be transcribed into multiple messenger RNAs (mRNAs), which over the course of hours are translated into hundreds to thousands of proteins. While many proteins are very stable with half-lives of hours to days, mRNAs tend to be degraded much more rapidly. In addition, these synthesis and degradation rates cover a wide range of values across different genes, explaining why the correlation between mRNA and protein concentrations in steady-state systems can be relatively low (Liu et al., 2016; Schwanhäusser et al., 2011; Vogel and Marcotte, 2012; McManus et al., 2015).

As technology has advanced, it is now possible to routinely estimate RNA and protein concentrations as well as the interactions between molecules for thousands of genes in parallel. As such we are ready to move beyond the analysis of single conditions, processes, or time points, to integrative work that compares multiple dimensions of gene regulation while assessing the dynamics of the system in time-resolved studies. First such analyses have emerged and revealed a picture of intricate coordination of RNA and protein synthesis and degradation. For example, during the response to lipopolysaccharide stimulation in dendritic cells, most gene expression changes are driven by transcription regulation (Jovanovic et al., 2015) – as is expected for this system that relies heavily on transcription signaling cascades (Amit et al., 2009; Bhatt et al., 2012; Garber et al., 2012).

In contrast, responses to other stimuli, such as the Unfolded Protein Response (UPR), largely affect translation (Wek and Cavener, 2007). As the cell reacts to an overload of unfolded proteins in the endoplasmic reticulum (ER), the UPR reduces the rate of translation and attempts to restore homeostasis (Harding et al., 1999; Ron and Walter, 2007). However, ER stress and the UPR also reprogram transcription as well as RNA and protein degradation through an intricate network of events. Initially the ER stress sensor *PERK* halts translation through inhibitory phosphorylation of initiation factor *elF2a* and decreases the ER’s protein folding load (Harding et al., 1999, 2000).

Global translation suppression is accompanied by the activation of translation for important UPR regulators and the transcription of downstream stress-response genes. Indeed, *elF2a* phosphorylation induces translation of the transcription factor *ATF4* reduced abundance of the ternary complex allows ribosomes to bypass inhibitory upstream open reading frames (uORFs) in the mRNA’s 5’ untranslated region (UTR)(Vattem and Wek, 2004). Another example of a uORF-regulated gene is *GADD34 (PPP1R15A*)(Lee et al., 2009) whose protein product stimulates *elF2a* dephosphorylation and therefore re-activates translation while sensitizing cells to apoptosis (Brush et al., 2003). Some studies suggest that many more genes that are resistant to translation inhibition by *elF2a* phosphorylation, or even increase in translation (Baird et al., 2014; Cullinan et al., 2003; Guan et al., 2014; Maity et al., 2016; Ventoso et al., 2012).

Another major transcription factor, *XBP1*, is activated through non-canonical splicing by the *IRE1* endonuclease. *IRE1* also targets mRNAs for degradation, further relieving the ER from the burden of protein synthesis (Calfon et al., 2002; Hollien et al., 2009). A third UPR transcription factor, *ATF6*, is released from the ER membrane upon accumulation of misfolded proteins and activated through proteolytic cleavage in the Golgi (Haze et al., 1999). Finally, the proteasome degrades irreparably damaged and ubiquitinated proteins (Plemper and Wolf, 1999) – illustrating how indeed, over the course of several hours, the cell systematically responds to ER stress at all levels of gene expression regulation, via both synthesis and degradation of mRNAs and proteins.

This elaborate regulatory network illustrates the need for integrative, time-resolved assessment of the coordination of these different processes. Indeed, earlier studies have shown that time plays an important factor during ER stress and the decision between survival and apoptosis (Li et al., 2010). In contrast to the transcription-driven response to lipopolysaccharide (Jovanovic et al., 2015), both transcription and translation have substantial contributions to the ER stress response, but show different temporal patterns (Kershaw et al., 2015; Liu et al., 2017; Quirós et al., 2017; Cheng et al., 2015; Guan et al., 2014).

Here, we provide one of the most comprehensive assessments of the gene expression response to ER stress available to date. We collected replicate samples at four time points (0, 1, 4, and 8 hours) from human cervical cancer cells treated with tunicamycin. Using RNA-seq and mass spectrometry, we determined the complete RNA and protein concentrations for >7,000 genes in the core dataset and for >14,000 genes in the extended data. Further, using ribosome footprinting and protein-occupancy profiling, we mapped the binding of ribosomes and non-ribosomal proteins to mRNA, which informs on translation as well as several aspects of RNA processing, respectively. Finally, we applied the same technologies to cells treated with hydrogen peroxide to elicit oxidative stress. ER stress and the UPR are tightly linked to the oxidative stress response, largely due to reactive oxygen species (ROS) produced during protein folding in the ER (Malhotra and Kaufman, 2007). In addition, *elF2a* phosphorylation and translation inhibition are part of Integrated Stress Response, which is conserved across eukaryotes and triggered by a variety of stresses (Dickhout et al., 2012; Harding et al., 2003; Pakos-Zebrucka et al., 2016). As the data rely on the coordinated change of two molecule types, e.g. a change in ribosome-bound mRNAs depends both on the number of translating ribosomes as well as the mRNA abundance, we have adapted our Protein Expression Control Analysis tool (PECA) (Teo et al., 2014, 2018) to deconvolute the measurements and extract significant regulatory events per gene, per time point, and per regulatory level. Therefore, the presented data explores both shared and stress-specific regulation, it describes both the early and later stress response, and it disentangles transcriptional from post-transcriptional regulation. It discusses general patterns seen across the core set of genes and specific examples that illustrate hypotheses on new pathways for the cell to respond to stress.

## Results

### Multiple data types describe gene expression regulation during stress

We present a resource comprising four complementary data types collected for cervical cancer cells: RNA and protein concentrations measured with RNA-seq and quantitative proteomics, respectively, and the binding profiles of ribosomes and non-ribosomal proteins along mRNAs measured with ribosome footprinting and protein occupancy profiling (Figure 1A). The data aims at providing a comprehensive picture of the different biological processes involved in gene expression regulation in response to stress, with a specific focus on post-transcriptional regulation during the Unfolded Protein Response. While ribosome footprints mapping to coding regions serve as an estimate of translation efficiency, footprints of non-ribosomal proteins mapping to the untranslated regions (UTRs) suggest the binding of other regulators of translation, RNA stability, and localization.

**Figure 1.**
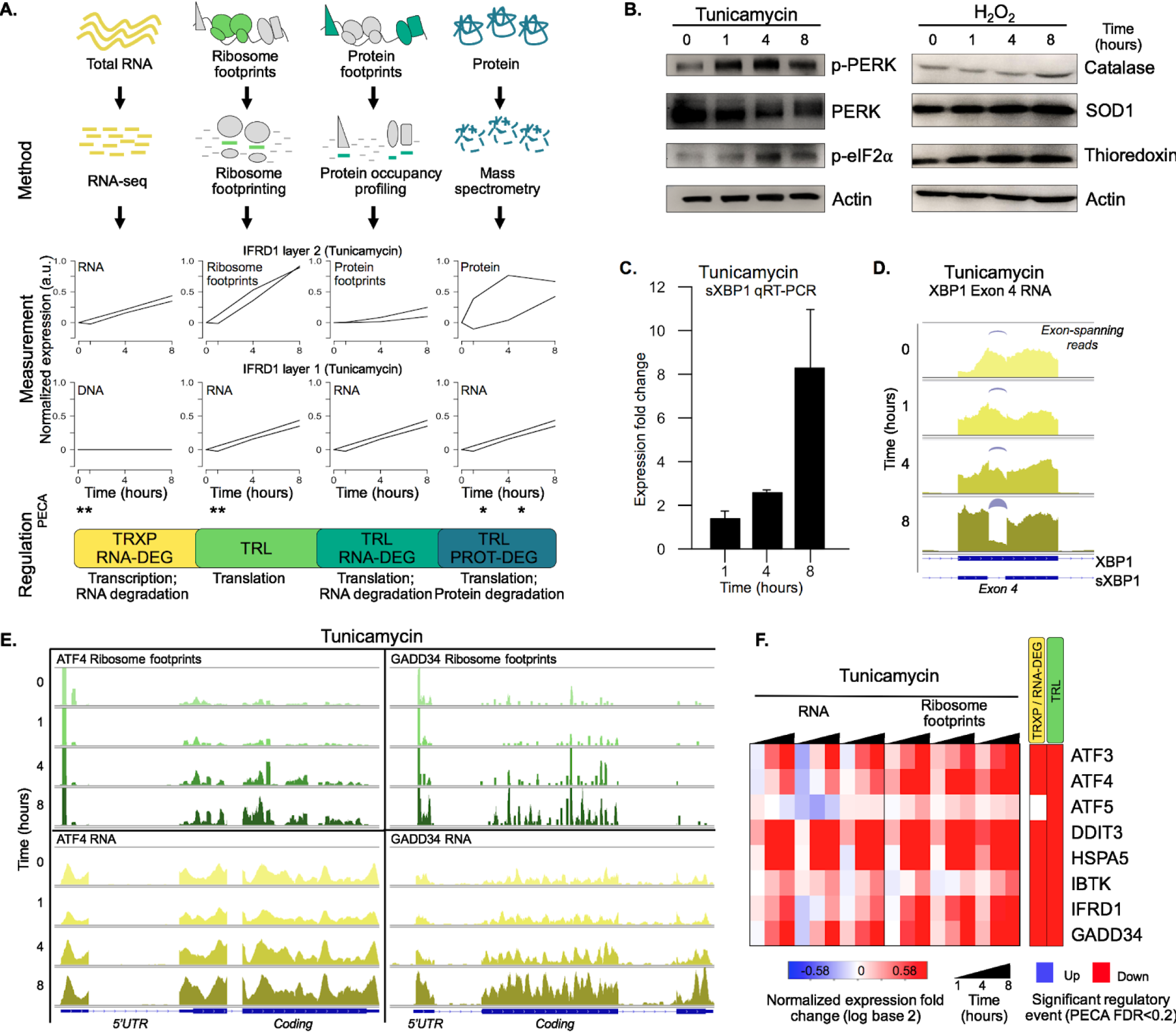
A multi-layered datasets describe the stress response. **A.** The schematic illustrates the experimental design that maps multiple layers of regulation in response to stress. RNA and protein abundances were measured using RNA-seq and mass spectrometry, respectively. Ribosome footprinting and protein occupancy profiling were used to map the binding of ribosomes and non-ribosomal proteins along mRNAs, respectively. Time points and genes with significant regulation were extracted from each data type with Protein Expression Control Analysis (PECA)(Teo et al., 2014, 2018). For changes in RNA concentration, assuming DNA concentration remains constant, PECA reports on transcription and RNA degradation regulation. After factoring out changes that occur in RNA concentrations, the ribosome footprinting, protein occupancy profiling, and mass spectrometry datasets inform on translation, binding of post-transcriptional regulations, and translation/protein degradation changes, respectively. The figure shows the expression patterns for *IFRD1* as an example of significant regulatory events identified by PECA; layer 1 represents the changes in molecular abundance on which layer 2 levels partially depend (false discovery rate < 0.2 in both biological replicates). **B.** Phosphorylation levels of ER stress markers PERK and eIF2α increase after treatment with tunicamycin (left). Protein abundance increases for common markers of oxidative stress in hydrogen peroxide treated cells (right). **C.** Splicing of *XBP1* (*sXBP1*) in tunicamycin treated cells increases, represented as mean fold change and standard error of the mean compared to the 0 hour time point. **D.** Reads mapping to exon 4 of *XBP1* in the RNA seq data indicate an increase in the spliced isoform in response to tunicamycin. Spliced reads spanning the 26 nucleotide intron and corresponding to *sXBP1* are designated at each time point. Darker colors indicate later time points. **E.** The profiles show ribosome footprints (top) and RNA coverage (bottom) for two ER stress response genes, *ATF4* and *GADD34*, that are induced in their translation. The panels show data for one replicate after 0, 1, 4, or 8 hours of tunicamycin treatment. **F.** The heatmap shows normalized fold changes for all three replicates for RNA and ribosome footprints measurements of eight ER stress response genes regulated via upstream open reading frames. As the PECA assessment indicates, the genes are indeed translationally upregulated. p-PERK – phosphorylated PERK; p-eIF2α – phosphorylated eIF2α; TRXP – transcription; TRL – translation; RNA-DEG – RNA degradation; PROT-DEG – protein degradation.

To provide a robust dataset that describes the dynamic nature of the stress response, we collected replicates for all data types across four time points (0, 1, 4, 8 hours). Further, we repeated the experiment subjecting the cells to oxidative stress enabling the identification of genes involved in the shared stress response and those specific to the UPR. From the extended sequencing data comprising >14,000 genes with triplicate measurements and >10,000 quantified proteins (**Tables S1-S4**), we compiled a core set of 7,011 genes with complete, replicate measurements (Figure 2, **Table S5**). We present general patterns and specific examples of regulation that we extracted from this core set.

**Figure 2.**
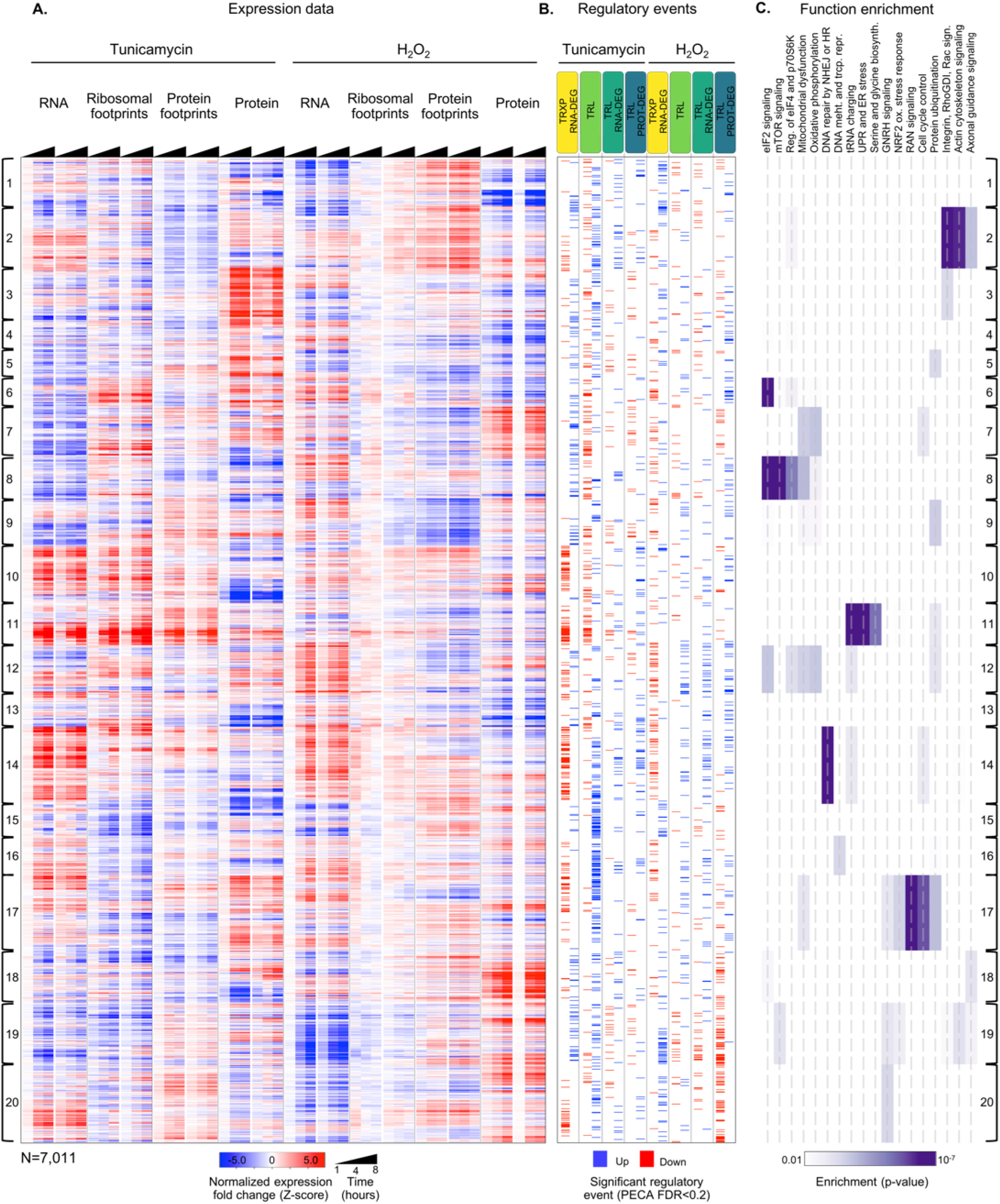
Integrated data reveal global and gene-specific regulation in response to stress. **A.** The heatmap shows relative changes in RNA abundance, ribosome and non-ribosomal protein footprints, and protein abundance for 7,011 genes (rows) in duplicate after 1, 4 and 8 hours of tunicamycin or hydrogen peroxide treatment. Rows are sorted using complete-linkage hierarchical clustering and labeled according to their cluster number. **B.** The heatmap indicates significant regulatory events for the genes at each regulatory level displayed along the top (false discovery rate < 0.2 for both replicates). **C.** The heatmap shows function pathways that are significantly enriched in the gene clusters (p-value < 0.01). The order of the genes is the same across all three panels. FDR – false discovery rate; TRXP – transcription; TRL – translation; RNA-DEG – RNA degradation; PROT-DEG – protein degradation

As expression changes are interdependent, for example protein expression depends on RNA changes, we apply our tool for Protein Expression Control Analysis (PECA)(Teo et al., 2014, 2018) to extract significant regulatory events at specific levels (Figure 1A). PECA is designed to ‘subtract’ the change in transcript abundance for each gene in order to isolate the contribution of translation and degradation to protein expression changes. PECA also takes into account the temporal information to report both significantly regulated genes *and* time points. Here we expand the use of PECA to all four datasets and convert concentration and binding data into information on regulatory events. For the RNA data, assuming DNA copy number does not change, PECA informs on significant changes in transcription and RNA degradation (TRXP/RNA-DEG) (Figure 1A). For ribosome footprinting data, PECA extracts information on translation (TRL) by controlling for changes at the RNA level. For the protein occupancy data, PECA extracts information on the binding of non-ribosomal regulators that can affect translation, RNA localization, and degradation (TRL/RNA-DEG). For the protein data, it extracts significant translation and protein degradation events (TRL/PROT-DEG). As such, PECA is capable of determining multiple levels of significant regulation per gene at each time point. To extract significant events, we apply the same threshold for the false discovery rate (FDR <0.2) for both replicates, however all results presented here are robust to the use of stricter cutoffs (**Figure** S1).

As an example, Figure 1A shows the observed data and the inferred significant regulatory events for the ER stress response gene *IFRD1* (indicated by asterisks). This gene is upregulated at the transcript level through reduced degradation during tunicamycin treatment, however it is also translationally induced via uORFs in its 5’ UTR, resulting in increased protein not solely due to mRNA abundance (Zhao et al., 2010). We indeed report changes at the RNA and protein level along with a significant increase in translation.

Several lines of evidence illustrate the quality of the data and the appropriate induction of stress markers. As expected, tunicamycin elicits ER stress in the cells, which activates PERK leading to the subsequent phosphorylation and inhibition of eIF2α (Figure 1B). When examining a set of 143 known UPR genes, many of the genes peak in their response at four fours after treatment (**Figure S1**). In comparison, the expression increase of markers of oxidative stress such as catalase, SOD1, and thioredoxin is modest (Figure 1B) consistent with the chosen concentration of H202 being at the lower limit for what is sufficient to induce oxidative stress in cancer cells (Nakamura et al., 2003). As the main focus of this work is to provide a clear picture of the ER stress response, the overall-mild oxidative stress response provides a comparison for regulatory events observed. Finally, expression for many housekeeping genes remains largely unchanged, indicating cells are largely non-apoptotic (**Figure S1**).

The timeline of ER stress in our experiment is further confirmed by the correct activation of UPR regulators, *XBP1, ATF4*, and *GADD34* (*PPP1R15A*) (Figure 1C-E). The non-canonical splicing of the fourth exon in *XBP1* results in a frame-shifted transcript (sXBP1) that produces the XBP1 transcription factor; this is one of the earliest steps in the ER stress response (Uemura et al., 2009). While we observe some constituent sXBP1 even in unstressed cells, the spliced isoform dominates by eight hours after tunicamycin treatment (Figure 1C,D). The spliced isoform is present in both the RT-qPCR and RNA-sequencing data, supporting our confidence in the resolution and quality of the large-scale dataset. We observe clear translation upregulation of *ATF4* and *GADD34* through an increase in ribosome footprints in the main open reading frame (Figure 1E). Both *ATF4* and *GADD34* are known to escape global translation shutdown and are translationally upregulated via uORFs in the 5’ UTR (Lee et al., 2009). *GADD34* is responsible for the dephosphorylation of eIF2α, making it essential for the reinitiation of translation necessary for cell survival and eventual apoptosis during chronic stress conditions (Adler et al., 1999; Hollander et al., 1997; Marciniak et al., 2004).

We validate PECA’s ability to extract significant regulatory events through examination of eight genes with known uORF-medidated translation increase during ER stress (Figure 1F). The genes include *ATF4, GADD34*, and *IFRD1* discussed above, as well as *ATF3, ATF5, DDIT3* (CHOP), *HSPA5* (BiP), and *IBTK*. Across all three replicates, PECA correctly identifies significant translation regulation based on the increase in ribosome footprints compared to RNA levels (TRL, Figure 1F). Wth the exception of *ATF5*, all genes also show concordant increases in RNA concentrations that are marked as significant regulatory events (TRXP/RNA-DEG). **Figure S1** shows additional uORF– and IRES-regulated genes that serve as a negative control: these genes are not associated with ER stress and PECA does not identify translation induction. **Figure S2** shows that the replicates are highly consistent.

### Shared and stress-specific regulatory patterns emerge

Figure 2 illustrates the complexity of the response to tunicamycin and hydrogen peroxide treatment and reveals multiple routes of gene expression regulation. As expected, translation repression upon phosphorylation of eIF2α is the most frequent response to ER stress, affecting approximately one sixth of the 7,11 core genes (Table 1, N(TRL down) = 1,189). However, many genes also increase in translation during ER stress (Table 1, N(TRL up) = 746), including those with known mechanisms of translation induction via uORFs, as mentioned above. ER stress also elicits a large response at the transcript level, and in contrast to translation, we find twice as many transcriptionally up– than down-regulated genes (Table 1, N(TRXP; RNA-DEG up) = 1,012 and N(TRXP; RNA-DEG down) = 590). In comparison, oxidative stress affects translation very little in our experiment (Table 1). PECA results for the extended data show similar regulatory distributions at each level (**Tables S1-S4**) and are robust to different significance thresholds (**Figure S1B**).

**Table 1.** Many genes are regulated at different levels during stress. The table summarizes the numbers of genes with significant regulatory events as defined by the PECA analysis of two paired data types (level 1 and level 2 as explained in Figure 1, false discovery rate < 0.2 in both replicates). The events are split into stress-specific and shared events. The shared events might be part of the Integrated Stress Response and show some significant function enrichment (false discovery rate<0.01, NCBI DAVID function annotation tool). All abbreviations are as defined in Figure 1. * – Overlap significant with p-value < 0.01 (hypergeometric test)

| Data type (level 2) | Data type (Regulation) |  | Specific to ER stress (tunicamycin) | Specific to oxidative stress (H<sub>2</sub>O<sub>2</sub>) | Shared response | Function enrichment amongst shared genes |
| --- | --- | --- | --- | --- | --- | --- |
| RNA | Transcription; RNA degradation (TRXP; RNA-DEG) | Up | 889 | 481 | 123* | DNA repair |
|  |  | Down | 529 | 417 | 61* | - |
| Ribosome footprints | Translation (TRL) | Up | 706 | 204 | 40* | - |
|  |  | Down | 1,116 | 249 | 73* | Cytoplasmic chaperones |
| Protein footprints | Translation; RNA degradation (TRL; RNA-DEG) | Up | 171 | 274 | 16* | - |
|  |  | Down | 371 | 359 | 40* | Cell adhesion |
| Protein | Translation; protein degradation (TRL; PROT-DEG) | Up | 235 | 614 | 24 | - |
|  |  | Down | 358 | 492 | 32 | - |

Confirming successful induction of the ER stress response, a number of genes with similar expression changes in response to tunicamycin is enriched in UPR pathways (Figure 2, cluster 11, p-value<0.0001) and a quarter of the genes are upregulated in both transcription and translation (71 of 302 genes). This cluster also encompasses aminoacyl-tRNA synthetases (p-value<2.6×10^-9) and serine biosynthesis enzymes (p-value<1,3×10^-6) that are regulated in a manner similar to the UPR genes and are discussed later.

We observe significant increase in protein expression during ER stress for a set of genes significantly enriched in the integrin signalling pathway (Figure 2, cluster 3, p<0.0001, **Figure S4**). Integrins are transmembrane proteins that are synthesized and processed in the ER (Tiwari et al., 2011), and indeed, cluster 3 is generally enriched in transmembrane proteins (Figure 2, 161 of 377 genes, p < 0.0001). Many of the genes decrease in translation during ER stress, but show stable or increasing protein abundance (**Figure S4**), suggesting that as the ER becomes incapable of synthesizing new transmembrane proteins during stress, those already present within membranes may exert increased stability to preserve cellular function until synthesis of new proteins is able to resume.

In our experiment, only about a tenth of significant regulatory events in each category is shared across the two stresses, as defined by genes being regulated in the same direction (Table 1, shared/specific genes ranging from 8 to 16% of total events). For most of the categories, the number of shared genes is significant, with the exception of translation and protein degradation (p-value < 0.05, hypergeometric test). In contrast to ER-stress-specific events which are dominated by translation, the shared response is dominated by up– and down-regulation of transcription and RNA degradation (Table 1, TRL vs. TRXP; RNA-DEG).

### Transcriptional and post-transcriptional changes can have the same but often also opposing directions

The genes of the shared stress response, i.e. genes whose transcription is induced in response to both ER and oxidative stress, are enriched in DNA repair pathways (Table 1, p-value < **0.01**). Many of these genes belong to cluster 14 in Figure 2A, and some are shown Figure 3A. **Figure S6** shows the extended set of DNA repair enzymes, some of which do not respond to ER stress. Even if the response is not significant with respect to our chosen cutoffs, many genes still increase in transcript levels (Figure 3A).

**Figure 3.**
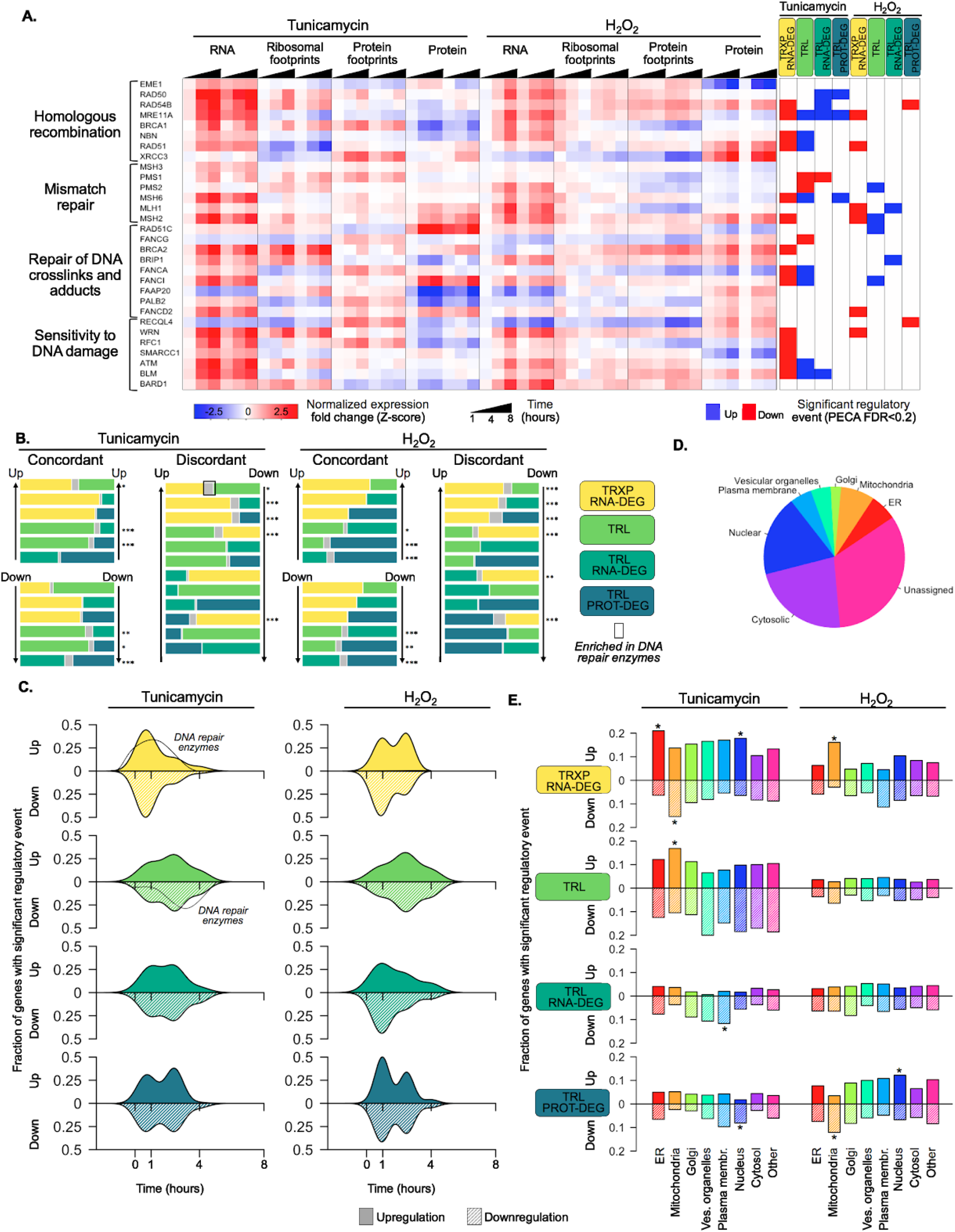
Gene expression is differentially regulated across levels, time, and organelles. **A.** The heatmap shows the relative changes in RNA abundance, ribosome and non-ribosomal protein footprints, and protein abundance for genes involved in selected DNA repair pathways, with significant regulation at each regulatory level as determined by PECA (false discovery rate < 0.2 in both replicates). **Figure S6** shows the data for all DNA repair genes. **B.** The venn diagrams illustrate the overlap (intersection) between genes regulated at different levels. Concordant or discordant regulation can be extracted through comparison of events affecting gene expression in the same or different directions, respectively. The color of the regulation type is encoded in the legend on the right. Bar sizes represent relative number of genes within each regulatory group, and the intersection in grey. Significant overlap was determined by a hypergeometric test: * p-value <0.1,** p-value <1x10^-3,*** p-value <1x10^-6. The box indicates group enriched in DNA repair genes. **C.** Smoothed density plots indicate the time-dependent occurrence of regulatory events. A positive y-axis represents upregulated genes, while negative y-axis represents downregulated genes. Traces indicate timing of transcription and translation of DNA repair genes significantly regulated during tunicamycin treatment. The panels show from top to bottom: TRXP/RNA-DEG; TRL; TRƯRNA-DEG; TRƯPROT-DEG. **D.** Pie chart illustrating the fraction of genes from the core dataset that encode proteins mapping to each organelle (Itzhak et al., 2016). **E.** Panels indicate the fraction of organelle-specific genes that are regulated at different levels, within each treatment. A positive y-axis represents upregulated genes, while negative y-axis represents downregulated genes. Fisher’s exact tests were used to assess whether up– or down-regulated genes were independent of each organelle. Significant differences in the distributions are indicated by a * (adjusted p-value<0.05). FDR – false discovery rate; TRXP – transcription; TRL – translation; RNA-DEG – RNA degradation; PROT-DEG – protein degradation

The DNA repair genes also illustrate how within one type of stress genes can be regulated in multiple ways. For example, RAD51, NBN, and MRE11A from the homologous recombination pathway are significantly induced in their transcript levels, but also significantly repressed in their translation in response to ER stress, counterbalancing the effect of transcription (Figure 3A, TRXP; RNA-DEG vs. TRL). Other examples include RAD54B, MLH1, and BLM, which increase in transcription, but decrease in their non-ribosomal protein binding or protein concentrations.

We examined these cases, in which genes are regulated in more than one way, more closely. In some cases, the events were discordant, such as shown in Figure 3A for the DNA repair genes: the cell appears to actively affect opposing processes. In other cases, the events were concordant, i.e. where the regulation followed the same direction, amplifying the effect. Figure 1F illustrates examples of such concordant regulation with coordinated transcription and translation for UPR genes like *ATF4, GADD34*, and *IFRD1*, and for genes in cluster 11 of **Figure 2**.

Indeed, both concordant and discordant regulation occurs within either of the stresses: for several categories, we find a significant number of genes regulated by two different categories, as indicated by the intersections (grey) between blocks in Figure 3B (hypergeometric test). Concordant regulation appears to be more frequent between the three categories that involve post-transcriptional processes (Figure 3B, TRL; TRL, RNA-DEG; TRL, PROT-DEG based on the ribosome and protein footprinting and protein concentration data, respectively). This concordance is perhaps due to translation being a common to the different categories. However, we also find mildly significant amount of concordant regulation between transcription and translation in response to ER stress, including many UPR genes mentioned above (Figure 3B, hypergeometric test, p-value <0.1; Figure 2, cluster 11).

Notably, a substantial number of genes is discordantly regulated in either stresses, and similar to the DNA repair genes discussed above and many genes in cluster 14 in Figure 2. Like for the DNA repair genes, discordance is commonly reflected by transcriptional and post-transcriptional regulation opposing each other (Figure 3B). Discordance can be implemented in different ways. For example, some genes, like MRE11A and NBN, show decreases in the abundance of translating ribosomes under ER stress and also a decrease in protein concentration, opposing the increase in mRNA concentration (Figure 3A). Other genes, like RAD50, BRCA1, and BRCA2, show an increase in both RNA abundance and ribosome footprints, however the increase in ribosome footprints is much smaller, reflecting an overall decrease in the translation rate for these transcripts which may or may not have consequences for protein concentrations.

Next, we examine the timing of the different regulatory processes. To do so, we averaged for each gene the time points at which significant regulation was observed across replicates and plot the smoothed frequency distributions (Figure 3C). The resulting traces for up– and down-regulation are surprisingly symmetrical. As an example, we also indicate the distribution for DNA repair genes. Their transcription is induced early after treatment, but translation is repressed throughout the mid-to-late phases of stress. This delay between the two events is perhaps due to a rapid increase in mRNA abundance upon the sensing of stress that subsequently outpaces the availability of ribosomes.

During ER stress, most of the transcript level changes occur early at one hour of tunicamycin treatment (Figure 3C). Some genes, such as those of the UPR, also change early in translation (Figure 1E, F). However, in general we observe a second translation response between one and four hours of treatment (Figure 3C). This bimodality is more pronounced for the protein expression data than for the ribosome footprinting data, and present in both ER and oxidative stress (Figure 3C). The delayed post-transcriptional response might reflect alternative mechanisms circumventing early translation shutdown or a recovery of protein synthesis from stress.

### Proteins from different subcellular localizations show different modes of regulation

The core set of 7,011 genes examined here localizes to both the cytosol and nucleus, as assigned from published data (Figure 3D)(Itzhak et al., 2016). Consistent with the above results, we observe much up-regulation of RNA levels (TRXP/RNA-DEG) upon tunicamycin treatment for genes whose proteins reside in the ER (adjusted p<0.009; Figure 3E). These genes include many known UPR targets that relieve the burden of accumulating misfolded proteins, but also the DNA repair genes discussed earlier.

Proteins localized to plasma membranes are marked by significantly reduced binding of non-ribosomal proteins to the 3’ UTR of the corresponding mRNAs (TRL/RNA-DEG; adjusted p-value<0.002; Figure 3E). Upon closer examination, we find that many of these genes include transmembrane proteins that require processing in the ER and therefore particularly burden the organelle during stress. These proteins often undergo N-linked glycosylation (p-value < 0.0001), the precise process blocked by tunicamycin and whose failure leads to misfolding. Therefore, it is tempting to speculate that the reduction in binding of non-ribosomal proteins to the mRNAs of these genes acts to repress the translation of these problematic mRNAs or trigger their degradation. When examining these transmembrane proteins more closely, we noticed that the former explanation is more likely than the latter: transcript levels of the proteins do not change or increase indicating stabilization of the mRNA, while ribosome binding decreases indicating reduced translation (**Figure S4**).

Proteins localized to mitochondria encompass the only category of genes which are significantly upregulated in translation (TRL, adjusted p-value<0.0003; Figure 3D). A similar, although non-significant trend is also observed at the protein level (TRL/PROT-DEG). Figure 4 and the section below discusses these genes in more detail.

**Figure 4.**
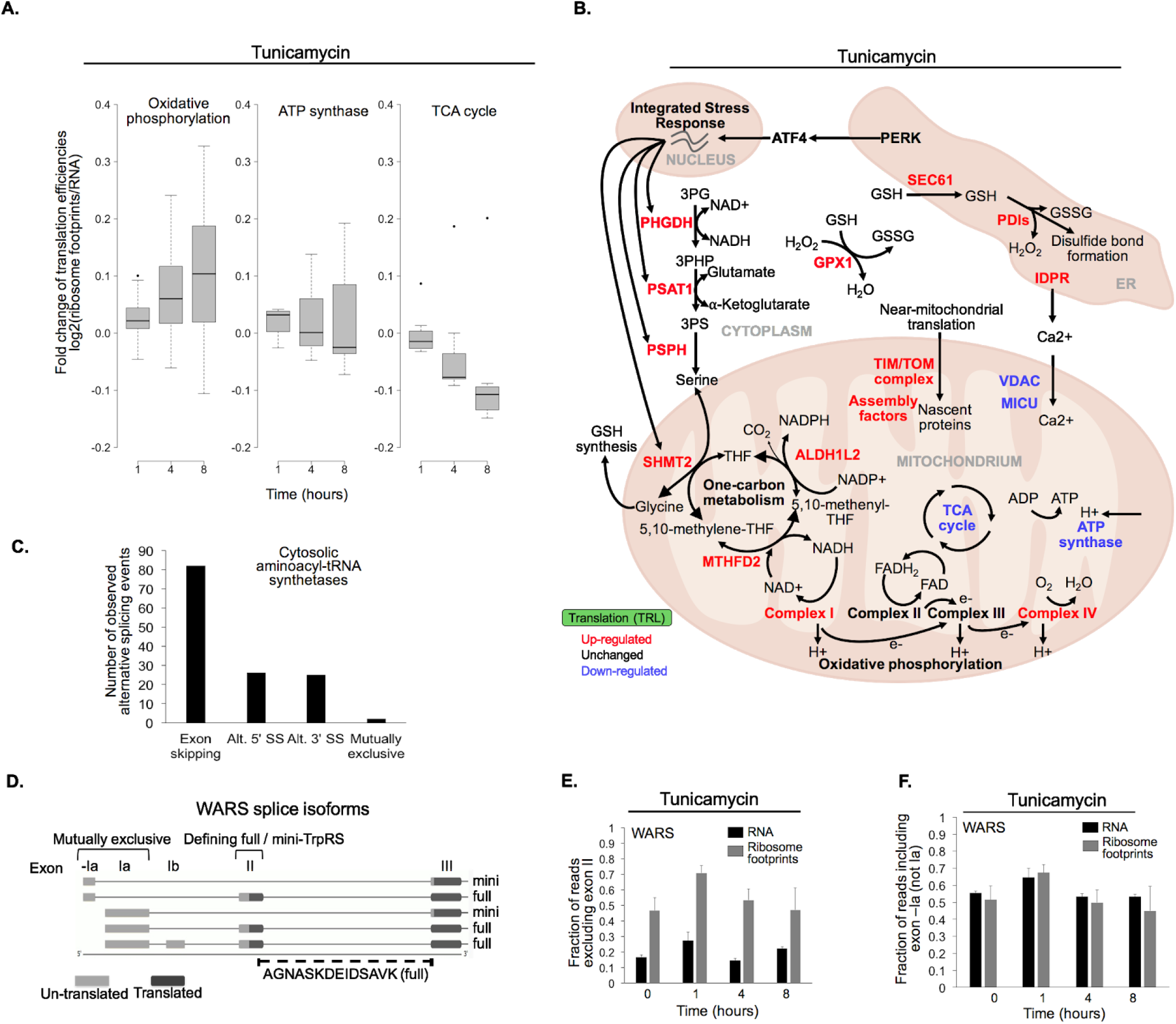
The combination of multiple datasets highlights new aspects of the stress response. **A.** The panels show the time-dependent changes in translation efficiency for different genes responding to tunicamycin treatment. The panels show the log base 2 ratio of ribosome footprints by RNA abundance as a proxy of translation efficiency, averaged over all genes and replicates. **Figure S7** shows the corresponding expression data for the respective genes. **B.** The diagram illustrates the simplified relationships between genes in panel **A.** Translation up– or down-regulation of genes and pathways is indicated by color. **C.** In our alternative splicing analysis of cytosolic aminoacyl-tRNA synthetases, we detected a total of 82 exon skipping events, 26 alternative 5’ splice sites (Alt. 5’ SS), 25 alternative 3’ splice sites (Alt. 3’ SS), and 2 mutually exclusive exons. **D.** The two protein isoforms of tryptophanyl-tRNA-synthetase (TrpRS, WARS) are encoded by five different transcript variants, each with a different 5’UTR that arises through alternative splicing. When exon II is spliced in, translation starts on exon II, leading to a full-length protein isoform. When exon II is skipped, translation starts on exon III, resulting in the truncated mini-TrpRS isoform. The peptide AGNASKDEIDSAVK spans the exon II/III junction and is only present in the full-length isoform. Its expression is shown in **Figure S5. E**. Skipping of WARS’ exon II, which results in the mini-TrpRS isoform, increases after one hour of tunicamycin treatment in both the RNA and ribosome footprinting data. Expression of exon II decreases again after the first hour of ER stress.-F. WARS’ exons –la and la are mutually exclusive. The fraction of reads including exon –Ia increases at one hour after tunicamycin treatment similar to the skipping of exon II shown in panel **E.** The exon II skipping event is significantly more likely to occur when transcription starts on exon –Ia (p-value<0.01, **Figure S5**).

### Translation regulation modulates energy metabolism during ER stress

Closer examination of genes whose proteins localize to mitochondria indicates a shift in energy metabolism in response to ER stress from the tricarboxylic acid (TCA) cycle to one-carbon metabolism and oxidative phosphorylation. Translationally up-regulated genes are enriched in subunits of Complexes I-IV from the oxidative phosphorylation pathway (Figure 4A, B). While the genes’ mRNAs levels often decrease during ER stress, ribosome binding and therefore translation increases significantly (p-value < 0.0001, Figure 4A, B). In contrast, translation is largely unchanged among subunits of the ATP synthase and decreases among enzymes involved in the tricarboxylic acid (TCA) cycle with the exception of one outlier, *ACO2* (Figure 4A, B).

Examination of the 5’ UTR of the genes for Complex I-IV reveals a sequence element that might explain that their differential translation increase during ER stress. Translation of mitochondrial genes is often regulated by the TISU element, the Translation Initiator of Short 5’ UTR (Elfakess and Dikstein, 2008; Elfakess et al., 2011). The TISU element is also part of mTOR-sensitive mRNAs which in turn confers resistance to translation inhibition under energy deprivation (Gandin et al., 2016; Sinvani et al., 2015). Therefore, we hypothesize that TISU elements may play a role in the translation regulation of mitochondrial genes during ER stress, as this condition challenges the cell’s energy balance. Indeed, we find the TISU element (SAASAUGGCGGC) highly enriched among the translation initiation sites of Complex I-IV genes (E-value=1.7X10^-8), but not amongst genes for ATP synthase subunits or TCA cycle enzymes (E-value=0.03 and E-value=0.1, respectively).

We also observe increased translation for other mitochondrial components that are related to either oxidative phosphorylation or the ER stress response, including one-carbon metabolism, TIM/TOM complexes, respiratory-chain assembly factors, and calcium uptake (Figure 4B; **Figure S7**). Of particular interest is the subset of one-carbon metabolism genes whose proteins locate to mitochondria (*SHMT2, MTHFD2*, and *ALDH1L2*). While the TCA cycle generally supplies NADH that feeds into the oxidative phosphorylation pathway to generate ATP, production of mitochondrial NADH can also be achieved through one-carbon metabolism, particularly under conditions of stress (Ducker and Rabinowitz, 2017). Translation increase of one-carbon metabolism genes with simultaneous translation repression of TCA cycle genes, as we observe in our data (Figure 4A, B, **Figure S7**), suggests a rerouting of NADH production.

Our data also indicates an important role of serine metabolism in this process. NADH production via one-carbon metabolism is driven by serine, which can be generated through the upstream serine biosynthesis pathway in the cytosol (*PHGDH, PSAT1, PSPH*)(Figure 4B). While enzymes of serine biosynthesis are known to be transcriptionally induced in various stresses (Zhao et al., 2016; Zhou et al., 2017), we find additional upregulation of ribosome binding and therefore translation for these genes (Figure 4B; **Figure S7**). We also find such concordant increase in transcription and translation for the mitochondrial one-carbon metabolism enzymes, but not for their cytosolic homologs, supporting our interpretation.

Finally, components of the TIM/TOM complexes and respiratory-chain assembly factors, also exhibit discordant regulation during ER stress, similar to Complex I-IV: they decrease in mRNA abundance but increase in translation (Figure 4B, **Figure S7**). *TIMM17A* represents an exception as it significantly decreases at the protein level, consistent with previous findings (Rainbolt et al., 2013). Co-translational import and proper assembly of oxidative phosphorylation complexes is orchestrated by TIM/TOM and complex specific assembly factors, and dysregulation of this process leads to complex defects implicated in many human diseases (Calvo et al., 2010; Mick et al., 2012; Vogel et al., 2005; Weraarpachai et al., 2009; Heide et al., 2012; Mckenzie and Ryan, 2010). Therefore, this shared regulatory pattern may reflect a coordinated response between the oxidative phosphorylation complexes and their assembly machinery, which work together to adjust mitochondrial metabolism when the cell is burdened by misfolded proteins in the ER.

### Alternative splicing of aminoacyl-tRNA synthetases is one example of extensive regulation during stress

As a case study, we examined another group of genes in the same cluster as the UPR and serine biosynthesis genes which shows substantial transcription and translation upregulation: aminoacyl-tRNA synthetases (Figure 2, cluster 11, **Figure S4**). Substantial post-transcriptional regulation of these enzymes in response to stress has been observed before without an explanation (Cheng et al., 2015; Ventoso et al., 2012). As the enzymes are known for their additional functions beyond aminoacylation of tRNAs (Guo and Schimmel, 2013) and alternative splicing underlying such ‘moonlighting’ (Lo et al., 2014), we hypothesized that stress-dependent expression of alternative transcript variant might explain some of discrepancy that we observe in our data between changes in ribosome binding and the corresponding protein levels (**Figure S4**).

Indeed, we find evidence for ~80 alternative splicing events amongst the 20 cytosolic aminoacyl-tRNA synthetases where the RNA reads on exon-exon junctions indicate the presence of a second transcript variant (Figure 4C). The distribution of these events is similar to what has been observed across the entire human genome (Sammeth et al., 2008), with exon skipping being the most frequent splicing event. A small number of these alternative splicing events changes in response to ER or oxidative stress consistently across the three replicates; these examples indicate potential stress-dependent splicing. One such example is the tryptophanyl-tRNA synthetase *WARS* (**Figure D-F**). Other examples are shown in **Figure S5** and **Table S6**.

*WARS* has five transcript variants (Figure 4D). Two transcript variants exclude exon II, leading to the production of a shorter protein called mini-TrpRS which misses the N-terminal protein interaction domain (Tolstrup et al., 1995; Wakasugi et al., 2002). Mini-TrpRS is non-catalytic. For both the RNA and the ribosome footprint data, reads signifying skipping of exon II and production of mini-TrpRS increased at one hour after tunicamycin induced ER stress (Figure 4E). This trend is confirmed by a decrease in the abundance of the full-length protein as seen in the proteomics data (**Figure S5**). As mini-TrpRS is known for its anti-proliferative function and its role in angiogenesis (Wakasugi et al., 2002), the cell might produce the short variant to reduce proliferation early during stress.

Excitingly, our data describes another splice event for *WARS* in response to ER stress, letting us hypothesize on a possible regulatory mechanism affecting the production of mini-TrpRS. Exons –la and la in the 5’ UTR of *WARS* are mutually exclusive (Figure 4D), and we find a substantial increase in the inclusion of exon –Ia at one hour after stress treatment, correlation with the exclusion of exon II (Figure 4F). Indeed, when combining the data for the two splicing events, we observe a significant bias towards exclusion of exon II and production of mini-TrpRS if exon –la instead of la is used (p-value < 0.001, **Figure S5**). We hypothesize that this splice event in the UTR influences skipping of exon II.

### Ribosomes and post-transcriptional regulators bind to RNA secondary structures

Finally, we show that stress-dependent binding of proteins in our data to predicted secondary structures in the mRNA’s coding and untranslated regions delivers new hypotheses for mechanisms and regulators underlying post-transcriptional regulation (Figure 5). The importance of RNA secondary structures in the UTR has been increasingly recognized, especially in response to stimuli (Leppek et al., 2017; Mustoe et al., 2018; Wu and Bartel, 2017). Structures in the 5’ UTR can form Internal Ribosomal Entry Sites (IRES), which bind to ribosomes and allow translation in a cap-independent manner (Holcik et al., 1999; Macejak and Sarnow, 1991), and inhibitory elements that prevent the bypass of ribosomes leading to reduced translation (Xue and Barna, 2015; Xue et al., 2014). In the 3’ UTR, secondary structures can mitigate protein binding events that impact stability of mRNAs, translation efficiency, and mRNA localization (Chang et al., 2010; Mignone et al., 2002). As our dataset includes RNA footprinting data for both ribosomes and (non-ribosomal) proteins, we examined the relationship between protein binding and a set of predicted RNA secondary structures highly conserved across the eukaryotic kingdom (Parker et al., 2011).

**Figure 5.**
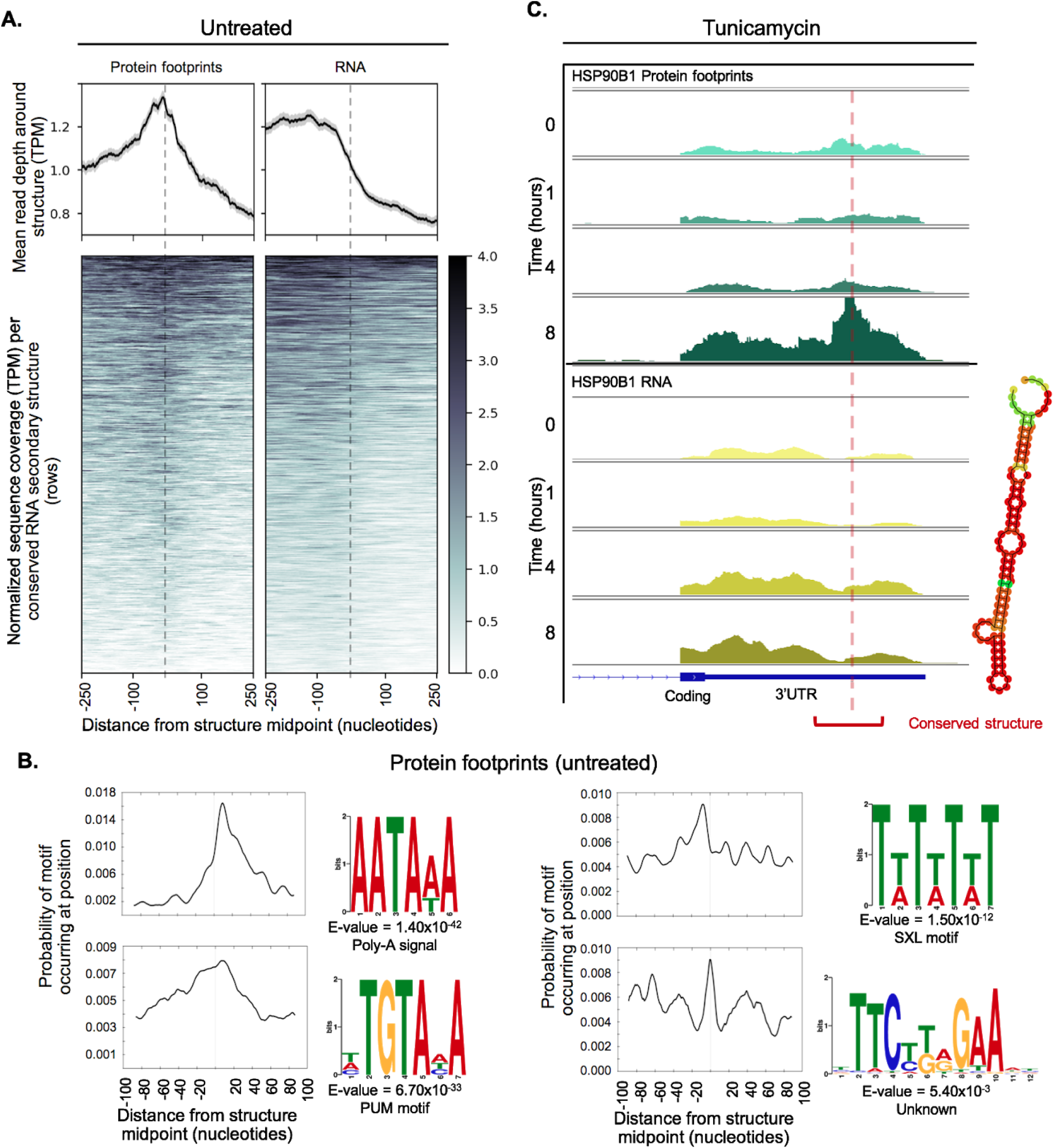
Conserved RNA secondary structures are enriched in protein footprints. **A.** Top panels show mean read depth (transcripts per million) of conserved RNA secondary structures in the **3’** UTR in a window +/- 250 nucleotides for protein footprints and RNA, respectively. Data was collected from untreated cells. Bottom panels show coverage for individual secondary structures over the 500 nucleotide window: each row corresponds to one secondary structure. Dashed lines indicate the midpoint of each secondary structure. **B.** The profiles show protein footprints (top) and RNA read coverage (bottom) for the **3’** UTR of the *HSP90B1* chaperone, for one replicate after 0, 1, 4, or 8 hours of tunicamycin treatment. Dashed lines indicates the midpoint of the predicted RNA secondary structure; the structure location is indicated by the red bracket. The predicted RNA secondary structure based on minimal free energy is shown on the right. **C.** The sequence logos illustrate motifs enriched in sequences corresponding protein-bound RNA secondary structures. Probabilities describe change of the motif occuring in a +/- 100 nucleotide window around the structure midpoints.

Indeed, the conserved secondary structures contain *bona fide* regulatory elements, as is demonstrated by recovery of known secondary structures, such as the iron response element. Iron response elements interact with ribosomes to regulate translation according to cellular iron levels (Aziz and Munro, 1987; Hentze et al., 1987a, 1987b). In total, we find transcript information for 302 secondary structures predicted in the genes’ 5’ UTR (Table 2, **Table S7**). Of these, two-thirds (190) have ribosome footprints mapping to a window 100 nucleotides upstream or downstream of the structure midpoint. Two RNA structures with highest abundance of ribosome footprints are located within 5’ UTR of the ferritin heavy and light chain transcripts, *FTH1* and *FTL* (**Figure S5E**). The structures are iron response elements, therefore validating the information content of the data.

**Table 2.** Conserved RNA secondary structures accumulate ribosome and non-ribosomal protein footprints. The table summarizes the combined analysis of conserved RNA secondary structures as predicted by reference (Parker et al., 2011) and the expression data. As expected, ribosome footprints accumulate around secondary structures in the 5’UTR of the genes more than in the 3’UTR, while this is the opposite for protein footprints. Listed numbers show data for all structures across the extended dataset (**Tables S1-S3);** the number of unique genes with predicted structures are indicated in parentheses. Supplemental Files

|  | 5' Untranslated region | 3' Untranslated region |
| --- | --- | --- |
| Conserved RNA secondary structures predicted in human genes | 760 (626) | 5,504 (2,789) |
| Conserved RNA secondary structures with RNA expression | 302 (277) | 2,697 (1,633) |
| Conserved RNA secondary structures with ribosome footprints | 190 (183) | 154 (151) |
| Conserved RNA secondary structures with protein footprints | 89 (81) | 1,808 (1,166) |

Post-transcriptional regulators binding to the 3’ UTR is a popular mode of expression control in our data. We observe a large number of genes with secondary structures predicted to the 3’ UTR, and for more than two thirds of them protein footprints map to a 200 nucleotide window around the structure midpoint (Table 2, 1,166 of 1,633 transcribed genes with structures). When averaging across all transcribed 3’ UTR structures in untreated data (0 hour time point), we find that protein footprints do in fact accumulate around these sites and are distributed symmetrically around the structure midpoint, suggesting proteins can directly bind these secondary structures (Figure 5A, **Table S7**). The RNA structures are predominantly located near the **3’** ends of transcripts, which results in declining read coverage downstream of the structures (Figure 5A, right panel, RNA).

The presence of sequence motifs further in close proximity to protein-bound structures further confirms the binding of regulatory factors, i.e. RNA-binding proteins (**Figure** 5B).The most significant motif corresponds to the poly(A) signal site (AAUAAA, E-value=1.40×10^-42). In our data this motif is most frequently 10 to 20 nucleotides downstream of protein bound RNA structures, in line with recent work showing *in vivo* folded 3’-end structures are often located near poly-A signals (Wu and Bartel, 2017)(Figure 5B).

Other observed motifs include predicted binding sites for Pumilio and Sex-Lethal (PUM and SXL), and a hairpin without any known RNA-binding partner (Figure 5B). Footprints for each of these motifs have a different spatial arrangement around the midpoint of structures. Pumilio RNA-binding sites display a broad distribution roughly centered around the structure (Figure 5B). SXL bindings sites have a distinct spacing pattern, with the highest frequency 8 to 10 nucleotides upstream of structure midpoints and additional sites roughly 10 to 15 nucleotides apart. Finally, the hairpin is often the center of the RNA structure itself, but also appears 30 to 40 nucleotides away from the structure (Figure 5B).

Finally, our data contains examples in which ER stress alters the occupancy of 5’ and 3’ UTR structured regions, either by ribosomes or non-ribosomal proteins, respectively. We identified the examples by comparing the local coverage of structures at 0 and 8 hour time points and identified significant changes (**Table S7**). We observe an increase in local ribosome footprints for a single 5’ UTR structure located within in the *TSC22D3* transcript (adjusted p-value = 0.031). This transcript encodes the glucocorticoid-induced leucine zipper protein (*GILZ*), that has been shown to act as a pro-survival factor during ER stress (André et al., 2016). We detect significant increases in local protein binding for five conserved 3’ UTR structures (adjusted p<0.01). Remarkably, these structures are all within genes that have an established role in the UPR, including three molecular chaperones that act as critical sensors of misfolded proteins: *HSP90B1, HSPA5*, and *CALR* (Lee, 2005; Huang et al., 2013; Eletto et al., 2010; Mungrue et al., 2009; Ellgaard and Helenius, 2003). We observe a clear increase in protein binding over these conserved secondary structures after eight hours of tunicamycin treatment – independent of changes in RNA reads mapping to the transcripts (Figure 5B,C, **Figure S5**).

## Discussion

### Both transcriptional and post-transcriptional regulation govern the stress response

We present a high-quality, information-rich resource that demonstrates the multiple levels at which cancer cells respond to either ER or oxidative stress. In capturing not only RNA and protein abundances, but also ribosome and protein footprints along mRNAs, we describe the highly dynamic character of various stress response pathways during the first eight hours of stress (Figure 2). Expanding our statistical tool PECA (Teo et al., 2014, 2018), we extract hundreds of genes with significant changes in transcription, translation, and degradation at individual time points. Our data is directed at improving our understanding of the response to ER stress, and provides novel insights into the complex nature of the Unfolded Protein Response not only by the global trends we identify, but also through hypothesis-driven analyses of specific pathways, e.g. stress specific splicing of aminoacyl-tRNA synthetases and the link between translation regulation and energy metabolism.

Translation regulation is a key component of the mammalian UPR, as translational shutdown is one of the earliest reactions to ER stress. However the extent to which genes can evade this global inhibition and the ways in which they do is an active area of investigation. Here we identify many genes that significantly increase in translation during ER stress (Table 1, Figure 2B), expanding upon the few well-known cases previously described (Vattem and Wek, 2004; Lee et al., 2009; Palam et al., 2011; Zhao et al., 2010; Zhou et al., 2008; Baird et al., 2014), as well as what has been suggested by polysome profiling experiments (Baird et al., 2014; Guan et al., 2017; Ventoso et al., 2012). Importantly, the increase in translation for many of these genes is independent of transcription. In fact, a majority are regulated through either transcription or translation, but not both (Figure 3B), as has been observed in other contexts (Tchourine et al., 2014).

Comparisons of our four data types reveals differences in the regulatory levels affecting gene expression. We observe that transcription regulation often occurs early in the stress response, while post-transcriptional regulation is bimodal, with both early and late changes (Figure 3C). All levels of regulation include genes that respond similarly in both stresses, comprising members of the shared stress response (Table 1). Within each stress, we observe both concordant and discordant regulation across different levels of regulation, i.e. processes that act in the same or opposite directions, respectively (Figure 3B). We find a statistically significant number of genes with discordance between their transcriptional and post-transcriptional regulation, which provides distinct patterns of expression for particular pathways during stress including DNA repair enzymes and mitochondrially localized proteins.

We present results that support the concept of an adaptive Unfolded Protein Response, where after surviving the acute phase of stress the cell adjusts via continual modulation of translation in order to recover protein synthesis and cope with the burdened ER (Guan et al., 2017; Imrie and Sadler, 2012; Urra et al., 2013). Indeed, we find that several key regulators of the UPR are concordantly upregulated in both transcription and translation (Figure 3B). Examining the timing of these events, we find that translation increases early after tunicamycin treatment and is followed later by an increase in transcription (Figure 1F, Figure 2) – suggesting that the cell adapts to longer stress exposure through stepwise amplification of the expression response.

In contrast, we hypothesize that transcriptional priming causes the discordant regulatory patterns that we observe for a different group of genes (Figure 3A-C). Many of these genes function in DNA damage repair, and they are transcriptionally induced in both stresses as part of the shared stress response. However, in the later phases of the ER stress response, the genes are unchanged or decrease in translation. Transcriptional priming has been well-established in plants where the transcriptome changes at the onset of stressful conditions and enables the fine-tuning of gene expression via translation later in the response (Conrath et al., 2015; Hilker et al., 2016). Indeed, translation decrease of DNA repair genes, likely through the PERƘ-mediated pathways (Oommen and Prise, 2013), is consistent with suppressed double-strand break repair during ER stress (Yamamori et al., 2013). As the UPR can switch from pro-survival to pro-apoptosis during persisting stress (Urra et al., 2013), the cell might initially promote DNA repair through transcriptional priming, but dial down this response until it either reaches new homeostasis or the decision to initiate cell death is made.

### Translation regulation links energy metabolism, mitochondria, and ER stress

To the best of our knowledge, our results offer one of first direct lines of support for translation, in addition to transcription, rerouting energy metabolism in response to ER stress (Leibovitch and Topisirovic, 2018; Pascal and Boiteau, 2011; 2015). We find a surprising and significant number genes localized to mitochondria that are upregulated in translation in response to ER stress (**Figure** 3E). Specifically, translation increases for genes involved in mitochondrial one-carbon metabolism (*SHMT2, MTHFD2, ALDH1L2*), but also in genes from serine biosynthesis (*PHGDH, PSAT1, PSPH*) which is upstream of one-carbon metabolism and localized to the cytosol (**Figure** 4A,B). Serine and one-carbon metabolism produce NADH amongst other products (**Figure** 4B)(Amelio et al., 2014). As serine biosynthesis diverts 3-phosphoglycerate from its use in glycolysis and the TCA cycle, the production of NADH via one-carbon metabolism may be used as an alternative to fuel Complex I and therefore drive oxidative phosphorylation, as has been suggested previously (Vazquez et al., 2011). Instead of using glycolysis and the TCA cycle for NADH production, the cell employs serine biosynthesis and one-carbon metabolism to generate ATP – and our results demonstrate that translation regulation supports this process during the response to ER stress (**Figure** 4A, B).

This resource reallocation from glycolysis to one-carbon metabolism is likely linked to the increased need for the reducing agent glutathione (GSH) during ER stress (Harding et al., 2003). GSH is essential in maintaining redox balance by supporting the formation of disulfide bonds in the ER and preventing the accumulation of reactive oxygen species (ROS)(Chakravarthi et al., 2006). Hydrogen peroxide and other ROS can be produced directly during the enormous efforts of the cell to refold proteins during ER stress (Zito et al., 2010; Guha et al., 2017). In fact we find that protein disulfide isomerases, main players in protein folding, are translationally induced during ER stress (Figure 4B). Their increased abundance will in turn heighten the demand for GSH in the ER to form disulfide bonds and in the cytosol to reduce the excess hydrogen peroxide.

Our data support several routes for the cell to accomplish this goal – mediated by translation (Figure 4A,B). One route is through increased translation of *SHMT2* which synthesizes glycine that is required for the biosynthesis of GSH (Amelio et al., 2014; Lu, 2013). Another route is through translation induction of subunits of the ER translocon *SEC61* that directs the uptake of GSH into the ER (Alder et al., 2005; Linxweiler et al., 2017; Ponsero et al., 2017). Finally, we observe translation induction of *GP×1*, which uses GSH to reduce hydrogen peroxide (Lubos et al., 2011).

Collectively, our data suggests the adjustment of translation to direct the flow of metabolism from glycolysis and the TCA cycle to serine biosynthesis and one-carbon metabolism during ER stress to not only support efficient synthesis of NADH for energy production by oxidative phosphorylation, but also to ensure GSH availability via glycine synthesis. Accordingly, we observe genes of the TCA cycle decrease in translation during ER stress, while genes of serine biosynthesis and one-carbon metabolism increase (Figure 4AB) – illustrating how translation links metabolism to the UPR.

### High-resolution data generate hypotheses on new regulators of the stress response and disease-relevant pathways

Our binding profile maps for ribosomes and other *trans*-acting factors serves as an important resource to generate new hypotheses on mechanisms that underlie the regulation of expression observed. As an example, we show that mRNA regions in the 3’ UTR that fold into conserved secondary structures are preferentially occupied by proteins (Figure 5A). Some of these structures are enriched for binding motifs of well-characterized RNA-binding proteins, such as Pumilio, that frequently occur directly on the bound RNA structure or in the surrounding area (Figure 5B). Pumilio is a well characterized post-transcriptional regulator across different conditions (Kurisaki et al., 2009; Qiu et al., 2011; Weidmann et al., 2014). It is thought to repress translation and localize to stress granules (Morris et al., 2008; Namkoong et al., 2018) which may indeed play a role in the stress response described here.

We also observe stress-dependent protein-mRNA binding that coincides with induced translation, including for the chaperone *HSP90B1* (*GRP94*) that has multiple functions in the ER and stress response (Eletto et al., 2010). Our analysis identified significant translation upregulation for *HSP90B1* at eight hours after tunicamycin treatment (**Figure S5**). We find this result to coincide with increased protein binding to a conserved stem-loop structure in *HSP90B1*’s 3’ UTR (Figure 5C). Such translation increase correlating with changes in protein binding to 3’ UTR structures is also observed for the chaperones *HSPA5* and *CALR* (**Figure S5**). Interestingly, all three examples are are UPR regulators (Vervliet et al., 2012; Yoshida, 2007). Upon ER stress, the mRNAs for *HSP90B1, HSPA5*, and *CALR* are released from the ER and translated in the cytosol (Reid et al., 2014). For *HSP90B1*, it has been shown that its ER localization is independent of known signal sequences, with unknown additional regulators (Pyhtila et al., 2008). It is tempting to speculate that the protein binding we observe at these secondary structures might indeed represent such regulators that drive the translocation and translation induction of these genes during stress.

These examples underscore the importance of integrating information from multiple levels of regulation to ascertain a complete picture of the cellular response to stress. ER stress is associated with a wide range of human diseases, e.g. cancer, neurodegeneration, and liver disease (Lin et al., 2007). Indeed we find many genes involved in Alzheimer’s, Parkinson’s, and Huntington’s disease to be induced in their translation (**Figure S4**). Other pathways we investigate in more detail, including those from serine biosynthesis and one-carbon metabolism, have strong links to cancer growth and survival (Amelio et al., 2014; Maddocks et al., 2013; Mattaini et al., 2016; Yang and Vousden, 2016). Finally, the transcriptional priming that we discuss has direct implications in various pathologies: mild ER stress exposure can enhance the cell’s ability to respond to later, additional stress and protect against several severe disease phenotypes (Inagi et al., 2008; Vacaru et al., 2014; Hara et al., 2011; Mendes et al., 2009). The transcription induction of DNA repair genes soon after tunicamycin treatment might be one way for the cell to achieve such stress protection. As the UPR acts as an essential switch between cellular survival and death (Schönthal, 2012; Cubillos-Ruiz et al., 2017; Scheper and Hoozemans, 2015), our data and findings offer potential routes for developing new therapeutic strategies to promote survival among healthy cells that struggle to cope with such insults to the proteome, as well as drive apoptosis among tumorigenic cells known to hijack the UPR to promote their own growth. Taken together our results will support future analyses across a wide range of topics in the biomedical field.

## Experimental procedures

### Cell culture and treatment

All samples for the total RNA, ribosome footprinting, protein occupancy profiling, and proteomics were derived from cells grown in parallel, arising from the same passage number. Cells were split just before the experiment and grown in parallel under identical conditions. Due to required protocols, we then prepared the samples for the total RNA-seq and ribosome in together, while the samples for the protein occupancy profiling and proteomics were processed separately. Biological replicates were collected independently on different days.

We grew Hela cells under standard condition, i.e in DMEM (Sigma) with 10% fetal bovine serum (Atlanta biologicals) and 1X penicillin streptomycin solution (Corning cellgro) at 37°C and 5% CO2. At ~60% confluency, we treated the cells with 60 μM H_2_O_2_ or 0.5 μg/ml tunicamycin to induce oxidative and ER stress, respectively. We treated samples 8, 4, 1, and 0 hours prior to collection and therefore collected all samples at the same time with similar confluency. For protein occupancy profiling, we added 200 μM of 4-thiouridine at ten hours before the treatment to incorporate photoreactive ribonucleoside analog required for protein occupancy profiling. For ribosome footprinting, we added 0.1 mg/ml cycloheximide for 5 min at 37°C before the harvesting the cells.

### Sample collection for total RNA and ribosome footprinting

All steps were performed according the the TruSeq Ribo Profile (Mammalian) kit protocol. Confluent plates of HeLa cells were first aspirated of their growth media and washed with fresh media supplemented with 0.1 mg/ml cycloheximide. The media was then removed and the cells were washed with 10 ml chilled PBS containing 0.1 mg/ml cycloheximide. Following removal of the PBS, 800 μl of Lysis Buffer was added and the cells were extensively scraped off plate and transferred to a pre-chilled eppendorf tube, recovering ~1 ml of lysate per sample. To insure complete lysis, we passed the lysate through a 25 gauge needle and further incubated on ice for ~10 min. The lysate was then clarified by centrifugation for 10 min at 20,000, 4°C and ~1 ml supernatant was recovered. For each treatment (tunicamycin and hydrogen peroxide), we aliquoted 100 μl of the cell lysate for total RNA extraction and 200ul for ribosome footprinting. The remaining lysate was frozen at -80C.

### Sample preparation

#### Total RNA profiling and ribosome footprinting

We used the TruSeq Ribo Profile (Mammalian) kit for both total RNA profiling and mapping of ribosome protected fragments (RPF). Briefly, for the total RNA samples, 100 μl aliquots of lysate were supplemented with 10 μl of 10% SDS and purified using a Zymo RNA Clean and Concentrator-25 Kits. For the RPF samples, we treated the cell lysate with 5 units of Truseq Ribo Profile Nuclease for each A260 of lysate and incubated for 45 mins at room temperature while agitating gently. We stopped reactions by adding SUPERase-In RNase inhibitor and placed the samples on ice. The nuclease-treated lysate (100 μl) was purified using Illustra Microspin S-400 columns as per instructions followed by addition of 10 μl of 10 % SDS to the flow-through. The remaining 100 μl of the nuclease digested RNA was stored in -80C for future use. Samples were purified using Zymo RNA Clean and Concentrator-25 kits and separated by denaturing 15% Urea-polyacrylamide gel electrophoresis (PAGE). We excised the desired size – corresponding to the ~28-30 nt range – using a dark field trans illuminator and purified according to the manufacturer’s protocol.

For library preparation, all samples were RiboZero treated and the total RNA samples were heat-fragmented at 94°C for 25 minutes according to the manufacturer’s protocol. Both the fragmented total RNA and RPF samples were then end-repaired and 3’ adapter ligated. Following adapter removal, the samples were reverse transcribed and the resulting cDNA was PAGE purified and circularized. We used one quarter of the circularized cDNA as a template for PCR amplification using Phusion (NEB) in a 50 μl volume using specific oligos (Truseq Riboprofile Forward PCR primer and Index PCR primer) and purified the samples with Agencount AMPure beads (Beckman Coulter). We purified the amplified libraries by using 8% Native PAGE and verified the final library size using the High Sensitivity DNA assay on the Agilent Bioanalyzer. We quantified samples using Qubit Fluorometric quantitation and sequenced them on a HiSeq 2500.

#### Protein occupancy profiling

After sample collection, cells were crosslinked with 365 nm UV light (0.2J/cm2) on ice using a Stratalinker 2400 (Stratagene). We scraped crosslinked cells off the plates with a rubber policeman, collected by centrifugation, washed with ice-cold PBS once and flash-froze the samples in liquid nitrogen for long-term storage.

Protein occupancy profiling was carried out as described previously (Munschauer, 2015). Briefly, we resuspended cell pellets in lysis/binding buffer (100 mM Tris-HCl pH 7.5 at 25 °C, 500 mM LiCl, 10 mM EDTA pH 8.0 at 25 °C, 1% LiDS, 5 mM DTT, Complete Mini EDTA-free protease inhibitor (Roche), incubated them at room temperature for 15 minutes for lysis and passed cells through a 21 gauge needle 10 times for shearing of genomic DNA. Lysates were incubated with oligo(dT) Dynabeads (Ambion) for 1 hr at room temperature on a rotating wheel. Following incubation, the beads were concentrated on a magnetic rack and the supernatant was stored on ice for further rounds poly(A)^+^-RNA depletion. We washed beads 3 times in lysis/binding buffer and 3 times in NP40 washing buffer (50 mM Tris-HCl pH 7.5 at 25 °C, 140 mM LiCl, 2 mM EDTA pH 8.0 at 25 °C, 0.5% NP40, 0.5 mM DTT) and crosslinked poly(A)^+^-RNA-protein complexes were eluted in low-salt elution buffer (10 mM Tris-HCI at at 25 °C) by incubation at 80°C for 2 minutes.

The stored supernatants were re-incubated with beads for 2 additional rounds of poly(A)^+^-RNA depletion following the described procedure. Eluates from different rounds of poly(A)^+^-RNA depletion are combined, incubated with RNase I for 10 minutes at 37°C and precipitated with 4 volumes of ammonium sulfate. The resuspended precipitate was separated by SDS-PAGE and transferred onto a nitrocellulose membrane. RNA-protein complexes were on-membrane incubated with Proteinase K for 30 minutes at 55°C to release protein-protected RNA fragments. RNA was recovered with phenol-chloroform extraction and subjected to small RNA library cloning procedure (Hafner et al., 2012). In short, RNA fragments generated by RNase I digestion were dephosphorylated with Calf intestinal alkaline phosphatase (CIP), radiolabeled at the 5’ end with [γ-^32^P]-ATP in a T4 Polynucleotide kinase (PNK) reaction followed by ligation of a pre-adenylated 3’ adapter and a 5’ adapter and reverse transcription to generate the cDNA library. In order to identify the RNA population with the desired fragment size, radiolabeled RNA size markers (24 and 50 nt) were used as ligation controls. The cDNA libraries were processed and subjected to sequencing on HiSeq 2500 following standard protocol (Hafner et al., 2012).

#### Proteomics analysis bv mass soectrometrv

For proteomics analysis, we resuspended cell pellets for each sample in 50 μl ice-cold PBS containing 1:100 protease inhibitor cocktail. The cells were then sonicated with probe sonicator 2 × 30 seconds with amplitude 5. The samples were returned to ice for 30 seconds between sonication intervals. After sonication, we mixed the samples with 50 μl trifluororethanol, and the mixtures were kept at 60 °C for 1 h. Then the samples were reduced in 15 mM DTT at 55 °C for 45 min, and alkylated in 55 mM iodoacetamide in the dark at room temperature for 30 min. Finally, we used 50 mM Tris (pH=8) to adjust the sample volume to 1 ml, and 1 ug mass spectrometry grade trypsin (Sigma Aldrich) was added to digest the proteins into peptides at 37 °C overnight.

We measured peptide concentrations with the Pierce Quantitative Fluorometric Peptide Assay (ThermoFisher) kit a. Tandem mass tag (TMT) 10-plex reagents (Thermo Scientific) were dissolved in anhydrous acetonitrile (0.8 mg/40 μl) according to manufacturer’s instruction. We labeled 30 ug/100 μl peptide per sample labelled with 41 μl of the TMT 10-plex label reagent at final acetonitrile concentration of 30% (v/v). Following incubation at room temperature for 1 h, we quenched the reactions with 8 μl of 5% hydroxylamine for 15 min. All samples were combined in a new microcentrifuge tubes at equal amounts and reduced to remove acetonitrile using an Eppendorf Concentrator Vacufuge Plus.

TMT-labelled tryptic peptides were subjected to high-pH reversed-phase high performance liquid chromatography fractionation using an Agilent 1200 Infinity Series with a phenomenex Kinetex 5u EVO C18 100A column (100 mm × 2.1 mm, 5 mm particle size). Mobile phase A was 20 mM ammonium formate, and B was 90% acetonitrile and 10% 20 mM ammonium formate. Both buffers were adjusted to pH 10. Peptides were resolved using a linear 120 min 0-40% acetonitrile gradient at a 100ul/min flow rate. Eluting peptides were collected in 2 min fractions. We combined about 70 fractions covering the peptide-rich region to obtain 40 samples for analysis. To preserve orthogonality, we combined fractions across the gradient, i.e. each of the concatenated samples comprising fractions which were 40 fractions apart. Re-combined fractions were reduced using an Eppendorf Concentrator Vacufuge Plus, desalted with C18 stage-tip, and suspended in 95% mass spectrometry grade water, 5% acetonitrile, and 0.1% formic acid for subsequent low pH chromatography and tandem mass spectrometry analysis.

For the first replicate, we used an EASY-nLC 1200 coupled on-line to a Fusion Lumos mass spectrometer (both Thermo Fisher Scientific). Buffer A (0.1% FA in water) and buffer B (0.1% FA in 80 % ACN) were used as mobile phases for gradient separation. A 75 μm × 15 cm chromatography column (ReproSil-Pur C18-AQ, 3 μm, Dr. Maisch GmbH, German) was packed in-house for peptides separation. Peptides were separated with a gradient of 5-40% buffer B over 110 min, 40%-100% B over 10 min at a flow rate of 300 nưmin. Full MS scans were acquired in the Orbitrap mass analyzer over a range of 300-1500 m/z at a resolution of 120,000 at m/z 200. The top 15 most abundant precursors were selected in data-dependent mode with an isolation window of 0.7 Thomsons and fragmented by higher-energy collisional dissociation with normalized collision energy of 40. MS/MS scans were acquired in the Orbitrap mass analyzer at a resolution of 30,000. The automatic gain control target value was 1e6 for full scans and 5e4 for MS/MS scans respectively, and the maximum ion injection time is 60 ms for both.

For the second replicate, we used an EASY-nLC 1000 coupled on-line to a Q Exactive spectrometer (both Thermo Fisher Scientific). Buffer A (0.1% FA in water) and buffer B (80% acetonitrile, 0.5% acetic acid) were used as mobile phases for gradient separation. An EASY Spray 50cm × 75μm ID PepMap C18 analytical HPLC column with 2μm bead size was used for peptide separation. We used a 110 minute linear gradient from 5% to 23% solvent B (80% acetonitrile, 0.5% acetic acid), followed by 20 minutes from 23% to 56% solvent B, and 10 minutes from 56% to 100% solvent B. Solvent B was held at 100% for another 10 minutes. Full MS scans were acquired with a resolution of 70,000, an AGC target of 1e6, with a maximum ion time of 120 ms, and scan range of 400 to 1500 m/z. Following each full MS scan, data-dependent high resolution HCD MS/MS spectra were acquired with a resolution of 35,000, AGC target of 1e5, maximum ion time of 250 ms, one microscan, 1.5 m/z isolation window, fixed first mass of 115 m/z, and NCE of 30.

### Validation experiments

#### αRT-PCR

For qPCR, we followed manufacturer’s instructions unless noted otherwise. We isolated total RNA from Hela cells treated with tunicamycin using Trizol extraction (Thermo Fisher Scientific) and purified RNA with the RNAesay mini Kit (Qiagen). We then synthesized cDNA using the Super Scriptl 11 First Strand cDNA synthesis kit (Invitrogen, Life technologies). We estimated *sXbp1* quantities by SYBR Green quantitative real-time PCR using Kapa Universal SYBR Green Supermix (Kapa biosystem) in a Roche Light Cycler 480 (Roche). All reactions were performed in triplicate. The expression of the spliced XBP1 was assessed by real time PCR (RT-PCR) in a Roche Light Cycler 480 (Roche) with KAPA universal SYBR green master mix. Relative gene expression was quantified using the ΔCT method with respective primers (*sXBP1* forward 5’-CTGAGTCCGCAGCAGGTG-3’; reverse 5’-GGGTCCAAGTTGTCCAGAATGC-3’) and normalized to *RPL19* (forward 5’-ATGTATCACAGCCTGTACCTG-3’; reverse 5’-TTCTTGGTCTCTTCCTCCTTG-3’). We used ΔΔCT method to determine the fold changes in the expression of sXbp1 (Livak and Schmittgen, 2001). Briefly, the threshold cycle (Ct) was determined and relative gene expression was calculated as follows: fold change = 2-Δ(ΔCt), where ΔCt (cycle difference) = Ct(target gene)-Ct(control gene) and Δ(ΔCt) = Ct(treated condition)-Ct(control condition).

#### Western Blotting

We boiled all samples collected at different time points after indicated treatment in sample buffer (Bio-Rad) supplemented with B-mercaptoethanol as per manufacturer’s instruction. We used equal amount of protein (25μg) from three independent grown cultures and treatments for blotting. The membrane was blocked using 5% BSA and incubated with respective antibodies (rabbit PERK (1:2000, CST C33E10), rabbit P-PERK (1:2000, CST 3179), rabbit P-eIF2alpha (1:2000, CST 9721S), rabbit anti Oxidative Stress Markers (Abcam ab179843). β-Actin (1:5000, CST 4967S) served as a loading control. We captured signal intensities of the bands in the Western blot with Kwikquant Imager (Kindle Biosciences, USA).

### Quantification and Statistical Analyses

#### RNA. ribosomal and (non-ribosomaΠ protein footprints

We used an in-house R pipeline to process the sequencing data from total RNA, ribosome footprinting and protein occupancy profiling. The fastq files of individual samples were processed with FASTX-Toolkit v0.0.14 to remove adapters and trim reads based on a minimum quality score of 20, as well as discard reads with a trimmed length shorter than 20 nucleotides. Reads mapping to ribosomal DNA were removed using Bowtie2 (Langmead and Salzberg, 2012). Remaining processed reads were aligned to the human reference genome (hg19) with TopHat v2.1.1 (Trapnell et al., 2012) using the gencode v19 GTF reference transcriptome (Harrow et al., 2012). Aligned bam files were filtered based on unique mapping and read length: RNA and protein footprints >25nt, ribosome footprints 28-30nt. Total counts per gene were calculated using HTseq (Planet et al., 2011). For expression measurements comprising the core dataset (Figure 2), RNA reads aligning to exons were counted as a measure of processed mRNA abundance, ribosomal footprint reads mapping to the coding regions of transcripts (CDS) were counted as a measure of translating ribosomes, and protein footprint reads mapping to the untranslated regions (UTR) were counted as a measure of protein bound UTR. Genes were filtered based on a minimum count of 10 across all samples and counts were normalized using the median ratio method implemented in DESeq2 (Anders and Huber, 2010; Love et al., 2014). Surrogate variables were estimated and removed via linear modeling (SVA) to remove batch effects (Leek and Storey, 2007). The data includes a total of ~14,000 genes and is described in **Tables S1-S3**.

#### Protein

We used MaxQuant Software version 1.5.5.1 with its integrated search engine Andromeda to analyse our raw files acquired from the mass spectrometer. Data was searched against the human sequence file (Homo_sapiens.GRCh37.75.pep.all.fa) sequence file downloaded from the ENSEMBL database (Cox and Mann, 2008; Cox et al., 2011; Tyanova et al., 2016). All sample fractions of two individual sets were grouped by setting up experimental design parameters in Maxquant. The mass tolerance of MS/MS spectra were set to 20 ppm with a posterior global FDR of 1% based on the reverse sequence of the human FASTA file. In addition, MS/MS data were searched by Andromeda for potential common mass spectrometry contaminants. Trypsin/P specificity was used to perform database searches, allowing two missed cleavages. Carbamidomethylation of cysteine residues and 10-plex TMT modifications on Lys and N-terminal amines were considered as a fixed modification, while oxidation of methionines and N-terminal acetylation were considered as variable modifications. TMT quantification was performed at MS2 level with default mass tolerance and other parameters. We then used the reporter ion intensities as estimates for protein abundance. A total of 10,399 protein groups were identified, including 0.01% reverse sequences and contaminants. Protein groups with no measurement among either replicate were then removed, as well as those identified by only one peptide in either replicate. After filtering, for each protein the geometric mean was calculated across all samples within one stress and the intensities were divided by this mean. The median of these ratios over all proteins was used as a size factor to account for differences in global sequence coverage between samples, similar to library size normalization for RNA sequencing and footprinting experiments. SVA was also applied to remove batch effects as described above (Leek and Storey, 2007). The data describes a total of N=7,255 protein groups and is presented in **Table S4**.

#### The core dataset

To derive the core dataset of 7,011 genes (Figure 2, **Table S5**), we first mapped all data to common ENSG identifiers. If several genes or isoforms mapped to the same data, we used the major/most abundant isoform as the group’s identifier. The data was then filtered for completeness across all replicates and resulted in the 7,11 genes presented here as the core dataset. For each time series experiment, we calculated the log base 10 ratios of the measurement at time x compared to the measurement at time 0. We then normalized across the entire time course (but for each dataset separately) by subtracting the average value from each entry and dividing by the standard deviation.

We visualized and analyzed the data matrix in PERSEUS version 1.5.5.1 (Tyanova and Cox, 2018), using hierarchical clustering, the ‘Correlation’ and ‘Complete’ options, marking values in blue-white-red scale (**Figure S3**). The principal components obtained from Principal Component Analysis (PCA) shows that all three replicates for the sequencing data are highly consistent and biologically meaningful (**Figure S3**). The data discussed in the main text focuses on two of the replicates and the first six eigenvectors of the subsequent PCA of the data. Using PERSEUS, we clustered the core dataset into 20 clusters with highest coherence in the similarity measure. The functional analyses were generated through the use of I PA (QIAGEN Inc.,https://www.qiagenbio-informatics.com/products/ingenuity-pathway-analysis), with an adjusted p-value cutoff of 0.0001.

#### Extracting significant regulatory events via Protein Expression Control Analysis (PECA)

To extract significant regulatory events, we adapted the statistical tool PECA that we developed and expanded recently (Teo et al., 2014, 2018). For PECA, we used the abundance data after SVA-based removal of surrogate variables, but prior to other transformations and normalization described above. The data was transformed by taking the natural logarithm and centered by subtracting the row median. We then smoothed the data using the Gaussian Process tool in PECAplus with standard settings and replicate information. We then performed the PECA analysis using standard settings as described in reference (Teo et al., 2014, 2018) for absolute, label-free data. In brief, PECA extracts significant regulatory events for each gene and each time point by examining the overall noise in the data and specifically the *change* in the synthesis and degradation of the respective molecule since and until the time points before and after the time point in question. For the footprinting data, synthesis and degradation are replaced by association and dissociation of the respective molecules.

The paired concentration data for the four different levels was derived as follows: i) for transcription and RNA degradation (TRXP; RNA-DEG) we paired the RNA abundance data with DNA abundance set to a constant; ii) for translation (TRL) we paired the ribosome footprinting with the RNA abundance data; iii) for translation and RNA degradation/localization/processing (TRL; RNA-DEG) we paired the protein footprinting with the RNA abundance data; and iv) for translation and protein degradation (TRL; PROT-DEG) we paired the protein and the RNA abundance data.

PECA reports for each gene a putative change point score. Given the overall score distribution it also calculates a false discovery rate (FDR) for each data point. We reported a significant regulatory event for a gene if we observed a change point with a score corresponding to an FDR<0.2 in both replicates, in the same direction (up or down), but regardless of the time point at which the event occurred. Stricter cutoffs resulted in very similar results (**Figure S1**). PECA results are provided in **Tables S1-S5**.

#### Alternative splicing analysis

We generated Sashimi plots (Katz et al., 2015) for the 20 cytosolic aminoacyl-tRNA synthetases on the Integrative Genomics Viewer (IGV) from RNA-seq data for all four time points and searched manually for alternative splicing events. These events were marked by reads that spanned exon-exon junctions and therefore unambiguously denoted inclusion or exclusion of specific exons. Single read events were excluded. To reduce the number of false positive splicing events, we only counted junction reads that started at or ended on at least one known exon boundary as determined by the RefSeq variants on IGV. Exons that were not annotated in these transcript variants were only included in the analysis if they comprised junction reads on both exon boundaries. We recorded the events as a major (V1) and minor (V2) splicing event (**Table S6**). We then examined the data for putative stress-dependent changes in production of these variants, requiring consistency across the three replicates.

#### Analysis of proteins binding to RNA secondary structures

We obtained a list of conserved RNA secondary structures from reference (Parker et al., 2011). Coverage of structures was calculated using htseq to count reads mapping within +/- 200 nucleotides of the midpoint. Structures were considered to be transcribed if they had an average coverage of 10 reads among untreated samples. Using R, we mapped these structures to the processed RNA, ribosome and protein footprinting data and evaluated the accumulation of reads around the structure (**Table S7**). We extracted motifs in the RNA sequence in and surrounding the secondary structure with the MEME package using standard settings (Bailey et al., 2006).

### Data availability

The data discussed in this publication have been deposited in NCBI’s Gene Expression Omnibus (Barrett et al., 2013; Edgar et al., 2002) and are accessible through GEO Series accession number GSE11317. The mass spectrometry data including the MaxQuant output files have been deposited to the ProteomeXchange Consortium via the PRIDE (Vizcaíno et al., 2016) partner repository with the dataset identifier PXD008575.

## Acknowledgements

The work was supported by the NIH/NIGMS grant 1R01GM113237-01 (to C.V.). We thank Kirsten Sadler-Edepli for providing the list of known UPR genes.

## Author contributions

J.R., Z.C., S.M., N.K., and S.K. performed experiments to generate data and validate results. J.R., Z.C., G.T., and B.P. processed and analyzed the data. J.R., S.M. M.M., K.M., Y.B.Z., and A.L. performed additional analyses. J.R, Z.C., S.M., N.K. K.A., M.L, L.F., S.K., H.C., and C.V. wrote the manuscript. All authors reviewed the manuscript. C.V. conceived the project and coordinated the effort.

---

Table S1. Extended RNA expression data

Table S2. Extended ribosome footprinting data

Table S3. Extended protein occupancy data

Table S4. Extended proteomics data

Table S5. Integrated core dataset of 7.011 genes

Table S6. Alternative splicing events across cytosolic aminoacvl-tRNA synthetases

Table S7. RNA secondary structures and ribosome/protein footprinting data

Supplemental Information. PDF with Figures S1-S7 and Supplemental Experimental Procedures.

## References

Adler, H.T., Chinery, R., Wu, D.Y., Kussick, S.J., Payne, J.M., Fornace, A.J., Jr, and Tkachuk, D.C. (1999). Leukemic HRX fusion proteins inhibit GADD34-induced apoptosis and associate with the GADD34 and hSNF5/INI1 proteins. Mol. Cell. Biol. 19, 7050–7060.

Alder, N.N., Shen, Y., Brodsky, J.L., Hendershot, L.M., and Johnson, A.E. (2005). The molecular mechanisms underlying BiP-mediated gating of the Sec61 translocon of the endoplasmic reticulum. J. Cell Biol. 168, 389–399.

Amelio, I., Cutruzzolá, F., Antonov, A., Agostini, M., and Melino, G. (2014). Serine and glycine metabolism in cancer. Trends Biochem. Sci. 39, 191–198.

Amit, I., Garber, M., Chevrier, N., Leite, A.P., Donner, Y., Eisenhaure, T., Guttman, M., Grenier, J.K., Li, W., Zuk, O., et al. (2009). Unbiased reconstruction of a mammalian transcriptional network mediating pathogen responses. Science 326, 257–263.

Anders, S., and Huber, W. (2010). Differential expression analysis for sequence count data. Genome Biol. 11, R106.

André, F., Corazao-Rozas, P., Idziorek, T., Quesnel, B., Kluza, J., and Marchetti, P. (2016). GILZ overexpression attenuates endoplasmic reticulum stress-mediated cell death via the activation of mitochondrial oxidative phosphorylation. Biochem. Biophys. Res. Commun. 478, 513–520.

Aziz, N., and Munro, H.N. (1987). Iron regulates ferritin mRNA translation through a segment of its 5’ untranslated region. Proceedings of the National Academy of Sciences 84, 8478–8482.

Bailey, T.L., Williams, N., Misleh, C., and Li, W.W. (2006). MEME: discovering and analyzing DNA and protein sequence motifs. Nucleic Acids Res. 34, W369–W373.

Baird, T.D., Palam, L.R., Fusakio, M.E., Willy, J.A., Davis, C.M., McClintick, J.N., Anthony, T.G., and Wek, R.C. (2014). Selective mRNA translation during eIF2 phosphorylation induces expression of IBTKα. Mol. Biol. Cell 25, 1686–1697.

Barrett, T., Wilhite, S.E., Ledoux, P., Evangelista, C., Kim, I.F., Tomashevsky, M., Marshall, K.A., Phillippy, K.H., Sherman, P.M., Holko, M., et al. (2013). NCBI GEO: archive for functional genomics data sets-update. Nucleic Acids Res. 41, D991–D995.

Bhatt, D.M., Pandya-Jones, A., Tong, A.-J., Barozzi, I., Lissner, M.M., Natoli, G., Black, D.L., and Smale, S.T. (2012). Transcript dynamics of proinflammatory genes revealed by sequence analysis of subcellular RNA fractions. Cell 150, 279–290.

Brush, M.H., Weiser, D.C., and Shenolikar, S. (2003). Growth arrest and DNA damage-inducible protein GADD34 targets protein phosphatase 1 alpha to the endoplasmic reticulum and promotes dephosphorylation of the alpha subunit of eukaryotic translation initiation factor 2. Mol. Cell. Biol. 23, 1292–1303.

Calfon, M., Zeng, H., Urano, F., Till, J.H., Hubbard, S.R., Harding, H.P., Clark, S.G., and Ron, D. (2002). IRE1 couples endoplasmic reticulum load to secretory capacity by processing the XBP-1 mRNA. Nature 415, 92–96.

Calvo, S.E., Tucker, E.J., Compton, A.G., Kirby, D.M., Crawford, G., Burtt, N.P., Rivas, M., Guiducci, C., Bruno, D.L., Goldberger, O.A., et al. (2010). High-throughput, pooled sequencing identifies mutations in NUBPL and FOXRED1 in human complex I deficiency. Nat. Genet. 42, 851–858.

Chakravarthi, S., Jessop, C.E., and Bulleid, N.J. (2006). The role of glutathione in disulphide bond formation and endoplasmic-reticulum-generated oxidative stress. EMBO Rep. 7, 271–275.

Chang, N., Yi, J., Guo, G., Liu, X., Shang, Y., Tong, T., Cui, Q., Zhan, M., Gorospe, M., and Wang, W. (2010). HuR uses AUF1 as a cofactor to promote p16INK4 mRNA decay. Mol. Cell. Biol. 30, 3875–3886.

Cheng, Z., Teo, G., Krueger, S., Rock, T., Koh, H.W.L., Choi, H., and Vogel, C. (2015). Differential dynamics of the mammalian mRNA and protein expression response to misfolding stress.

Conrath, U., Beckers, G.J.M., Langenbach, C.J.G., and Jaskiewicz, M.R. (2015). Priming for Enhanced Defense. Annu. Rev. Phytopathol. 53, 97–119.

Cox, J., and Mann, M. (2008). MaxQuant enables high peptide identification rates, individualized p.p.b.-range mass accuracies and proteome-wide protein quantification. Nat. Biotechnol. 26, 1367–1372.

Cox, J., Neuhauser, N., Michalski, A., Scheltema, R.A., Olsen, J.V., and Mann, M. (2011). Andromeda: a peptide search engine integrated into the MaxQuant environment. J. Proteome Res. 10, 1794–1805.

Cubillos-Ruiz, J.R., Bettigole, S.E., and Glimcher, L.H. (2017). Tumorigenic and Immunosuppressive Effects of Endoplasmic Reticulum Stress in Cancer. Cell 168, 692–706.

Cullinan, S.B., Zhang, D., Hannink, M., Arvisais, E., Kaufman, R.J., and Diehl, J.A. (2003). Nrf2 is a direct PERK substrate and effector of PERK-dependent cell survival. Mol. Cell. Biol. 23, 7198–7209.

Dickhout, J.G., Carlisle, R.E., Jerome, D.E., Mohammed-Ali, Z., Jiang, H., Yang, G., Mani, S., Garg, S.K., Banerjee, R., Kaufman, R.J., et al. (2012). Integrated stress response modulates cellular redox state via induction of cystathionine γ-lyase: cross-talk between integrated stress response and thiol metabolism. J. Biol. Chem. 287, 7603–7614.

Ducker, G.S., and Rabinowitz, J.D. (2017). One-Carbon Metabolism in Health and Disease. Cell Metab. 25, 27–42.

Edgar, R., Domrachev, M., and Lash, A.E. (2002). Gene Expression Omnibus: NCBI gene expression and hybridization array data repository. Nucleic Acids Res. 30, 207–210.

Eletto, D., Dersh, D., and Argon, Y. (2010). GRP94 in ER quality control and stress responses. Semin. Cell Dev. Biol. 21, 479–85.

Elfakess, R., and Dikstein, R. (2008). A Translation Initiation Element Specific to mRNAs with Very Short 5’UTR that Also Regulates Transcription. PLoS One 3, e3094.

Elfakess, R., Sinvani, H., Haimov, O., Svitkin, Y., Sonenberg, N., and Dikstein, R. (2011). Unique translation initiation of mRNAs-containing TISU element. Nucleic Acids Res. 39, 7598–7609.

Ellgaard, L., and Helenius, A. (2003). Quality control in the endoplasmic reticulum. Nat. Rev. Mol. Cell Biol. 4, 181–191.

Gandin, V., Masvidal, L., Hulea, L., Gravel, S.-P., Cargnello, M., McLaughlan, S., Cai, Y., Balanathan, P., Morita, M., Rajakumar, A., et al. (2016). nanoCAGE reveals 5’ UTR features that define specific modes of translation of functionally related MTOR-sensitive mRNAs. Genome Res. 26, 636–648.

Garber, M., Yosef, N., Goren, A., Raychowdhury, R., Thielke, A., Guttman, M., Robinson, J., Minie, B., Chevrier, N., Itzhaki, Z., et al. (2012). A high-throughput chromatin immunoprecipitation approach reveals principles of dynamic gene regulation in mammals. Mol. Cell 47, 810–822.

Guan, B.-J., Krokowski, D., Majumder, M., Schmotzer, C.L., Kimball, S.R., Merrick, W.C., Koromilas, A.E., and Hatzoglou, M. (2014). Translational control during endoplasmic reticulum stress beyond phosphorylation of the translation initiation factor eIF2α. J. Biol. Chem. 289, 12593–12611.

Guan, B.-J., van Hoef, V., Jobava, R., Elroy-Stein, O., Valasek, L.S., Cargnello, M., Gao, X.-H., Krokowski, D., Merrick, W.C., Kimball, S.R., et al. (2017). A Unique ISR Program Determines Cellular Responses to Chronic Stress. Mol. Cell 68, 885–900.e6.

Guha, P., Kaptan, E., Gade, P., Kalvakolanu, D.V., and Ahmed, H. (2017). Tunicamycin induced endoplasmic reticulum stress promotes apoptosis of prostate cancer cells by activating mTORC1. Oncotarget 8, 68191–68207.

Guo, M., and Schimmel, P. (2013). Essential nontranslational functions of tRNA synthetases. Nat. Chem. Biol. 9, 145–153.

Hafner, M., Renwick, N., Farazi, T.A., Mihailović, A., Pena, J.T.G., and Tuschl, T. (2012). Barcoded cDNA library preparation for small RNA profiling by next-generation sequencing. Methods 58, 164–170.

Hara, H., Kamiya, T., and Adachi, T. (2011). Endoplasmic reticulum stress inducers provide protection against 6-hydroxydopamine-induced cytotoxicity. Neurochem. Int. 58, 35–43.

Harding, H.P., Zhang, Y., and Ron, D. (1999). Protein translation and folding are coupled by an endoplasmic-reticulum-resident kinase. Nature 397, 271–274.

Harding, H.P., Zhang, Y., Bertolotti, A., Zeng, H., and Ron, D. (2000). Perk is essential for translational regulation and cell survival during the unfolded protein response. Mol. Cell 5, 897–904.

Harding, H.P., Zhang, Y., Zeng, H., Novoa, I., Lu, P.D., Calfon, M., Sadri, N., Yun, C., Popko, B., Paules, R., et al. (2003). An integrated stress response regulates amino acid metabolism and resistance to oxidative stress. Mol. Cell 11, 619–633.

Harper, J.W., Wade Harper, J., and Bennett, E.J. (2016). Proteome complexity and the forces that drive proteome imbalance. Nature 537, 328–338.

Harrow, J., Frankish, A., Gonzalez, J.M., Tapanari, E., Diekhans, M., Kokocinski, F., Aken, B.L., Barrell, D., Zadissa, A., Searle, S., et al. (2012). GENCODE: the reference human genome annotation for The ENCODE Project. Genome Res. 22, 1760–1774.

Haze, K., Yoshida, H., Yanagi, H., Yura, T., and Mori, K. (1999). Mammalian transcription factor ATF6 is synthesized as a transmembrane protein and activated by proteolysis in response to endoplasmic reticulum stress. Mol. Biol. Cell 10, 3787–3799.

Heide, H., Bleier, L., Steger, M., Ackermann, J., Dröse, S., Schwamb, B., Zörnig, M., Reichert, A.S., Koch, I., Wittig, I., et al. (2012). Complexome profiling identifies TMEM126B as a component of the mitochondrial complex I assembly complex. Cell Metab. 16, 538–549.

Hentze, M., Caughman, S., Rouault, T., Barriocanal, J., Dancis, A., Harford, J., and Klausner, R. (1987a). Identification of the iron-responsive element for the translational regulation of human ferritin mRNA. Science 238, 1570–1573.

Hentze, M.W., Rouault, T.A., Caughman, S.W., Dancis, A., Harford, J.B., and Klausner, R.D. (1987b). A cis-acting element is necessary and sufficient for translational regulation of human ferritin expression in response to iron. Proceedings of the National Academy of Sciences 84, 6730–6734.

Hilker, M., Schwachtje, J., Baier, M., Balazadeh, S., Bäurle, I., Geiselhardt, S., Hincha, D.K., Kunze, R., Mueller-Roeber, B., Rillig, M.C., et al. (2016). Priming and memory of stress responses in organisms lacking a nervous system. Biol. Rev. Camb. Philos. Soc. 91, 1118–1133.

Holcik, M., Lefebvre, C., Yeh, C., Chow, T., and Korneluk, R.G. (1999). A new internal-ribosome-entry-site motif potentiates XIAP-mediated cytoprotection. Nat. Cell Biol. 1, 190–192.

Hollander, M.C., Christine Hollander, M., Zhan, Q., Bae, I., and Fornace, A.J. (1997). MammalianGADD34, an Apoptosis– and DNA Damage-inducible Gene. J. Biol. Chem. 272, 13731–13737.

Hollien, J., Lin, J.H., Li, H., Stevens, N., Walter, P., and Weissman, J.S. (2009). Regulated Ire1-dependent decay of messenger RNAs in mammalian cells. J. Cell Biol. 186, 323–331.

Huang, C.-H., Chu, Y.-R., Ye, Y., and Chen, X. (2013). Role of HERP and a HERP-related Protein in HRD1-dependent Protein Degradation at the Endoplasmic Reticulum. J. Biol. Chem. 289, 4444–4454.

Imrie, D., and Sadler, K.C. (2012). Stress management: How the unfolded protein response impacts fatty liver disease. J. Hepatol. 57, 1147–1151.

Inagi, R., Kumagai, T., Nishi, H., Kawakami, T., Miyata, T., Fujita, T., and Nangaku, M. (2008). Preconditioning with endoplasmic reticulum stress ameliorates mesangioproliferative glomerulonephritis. J. Am. Soc. Nephrol. 19, 915–922.

Itzhak, D.N., Tyanova, S., Cox, J., and Borner, G.H. (2016). Global, quantitative and dynamic mapping of protein subcellular localization. Elife 5.

Jovanovic, M., Rooney, M.S., Mertins, P., Przybylski, D., Chevrier, N., Satija, R., Rodriguez, E.H., Fields, A.P., Schwartz, S., Raychowdhury, R., et al. (2015). Immunogenetics. Dynamic profiling of the protein life cycle in response to pathogens. Science 347, 1259038.

Katz, Y., Wang, E.T., Silterra, J., Schwartz, S., Wong, B., Thorvaldsdóttir, H., Robinson, J.T., Mesirov, J.P., Airoldi, E.M., and Burge, C.B. (2015). Quantitative visualization of alternative exon expression from RNA-seq data. Bioinformatics 31, 2400–2402.

Kershaw, C.J., Costello, J.L., Talavera, D., Rowe, W., Castelli, L.M., Sims, P.F.G., Grant, C.M., Ashe, M.P., Hubbard, S.J., and Pavitt, G.D. (2015). Integrated multi-omics analyses reveal the pleiotropic nature of the control of gene expression by Puf3p. Sci. Rep. 5, 15518.

Kurisaki, I., Watanabe, H., and Tanaka, S. (2009). Simulation Study of RNA-Binding Protein, Pumilio. J. Comput. Chem. Jpn. 8, 41–50.

Langmead, B., and Salzberg, S.L. (2012). Fast gapped-read alignment with Bowtie 2. Nat. Methods 9, 357–359.

Lee, A.S. (2005). The ER chaperone and signaling regulator GRP78/BiP as a monitor of endoplasmic reticulum stress. Methods 35, 373–381.

Lee, Y.-Y., Cevallos, R.C., and Jan, E. (2009). An upstream open reading frame regulates translation of GADD34 during cellular stresses that induce elF2alpha phosphorylation. J. Biol. Chem. 284, 6661–6673.

Leek, J.T., and Storey, J.D. (2007). Capturing heterogeneity in gene expression studies by surrogate variable analysis. PLoS Genet. 3, 1724–1735.

Leibovitch, M., and Topisirovic, I. (2018). Dysregulation of mRNA translation and energy metabolism in cancer. Adv. Biol. Regul. 67, 30–39.

Leppek, K., Das, R., and Barna, M. (2017). Functional 5’ UTR mRNA structures in eukaryotic translation regulation and howto find them. Nat. Rev. Mol. Cell Biol. 19, 158–174.

Li, H., Korennykh, A.V., Behrman, S.L., and Walter, P. (2010). Mammalian endoplasmic reticulum stress sensor IRE1 signals by dynamic clustering. Proc. Natl. Acad. Sci. U. S. A. 107, 16113–16118.

Lin, J.H., Walter, P., and Yen, T.S.B. (2007). Endoplasmic Reticulum Stress in Disease Pathogenesis. Annu. Rev. Pathol.: Mech. Dis. 0, 071003161323003.

Linxweiler, M., Schick, B., and Zimmermann, R. (2017). Let’s talk about Secs: Sec61, Sec62 and Sec63 in signal transduction, oncology and personalized medicine. Signal Transduction and Targeted Therapy 2, 17002.

Liu, T.-Y., Huang, H.H., Wheeler, D., Xu, Y., Wells, J.A., Song, Y.S., and Wiita, A.P. (2017). Time-Resolved Proteomics Extends Ribosome Profiling-Based Measurements of Protein Synthesis Dynamics. Cell Syst 4, 636–644. e9.

Liu, Y., Beyer, A., and Aebersold, R. (2016). On the Dependency of Cellular Protein Levels on mRNA Abundance. Cell 165, 535–550.

Livak, K.J., and Schmittgen, T.D. (2001). Analysis of Relative Gene Expression Data Using Real-Time Quantitative PCR and the 2-ΔΔCT Method. Methods 25, 402–408.

Lo, W.-S., Gardiner, E., Xu, Z., Lau, C.-F., Wang, F., Zhou, J.J., Mendlein, J.D., Nangle, L.A., Chiang, K.P., Yang, X.-L., et al. (2014). Human tRNA synthetase catalytic nulls with diverse functions. Science 345, 328–332.

Love, M.I., Huber, W., and Anders, S. (2014). Moderated estimation of fold change and dispersion for RNA-seq data with DESeq2. Genome Biol. 15, 550.

Lu, S.C. (2013). Glutathione synthesis. Biochim. Biophys. Acta 1830, 3143–3153.

Lubos, E., Loscalzo, J., and Handy, D.E. (2011). Glutathione peroxidase-1 in health and disease: from molecular mechanisms to therapeutic opportunities. Antioxid. Redox Signal. 15, 1957–1997.

Macejak, D.G., and Sarnow, P. (1991). Internal initiation of translation mediated by the 5’ leader of a cellular mRNA. Nature 353, 90–94.

Maddocks, O.D.K., Berkers, C.R., Mason, S.M., Zheng, L., Blyth, K., Gottlieb, E., and Vousden, K.H. (2013). Serine starvation induces stress and p53-dependent metabolic remodelling in cancer cells. Nature 493, 542–546.

Maity, S., Rajkumar, A., Matai, L., Bhat, A., Ghosh, A., Agam, G., Kaur, S., Bhatt, N.R., Mukhopadhyay, A., Sengupta, S., et al. (2016). Oxidative Homeostasis Regulates the Response to Reductive Endoplasmic Reticulum Stress through Translation Control. Cell Rep. 16, 851–865.

Malhotra, J.D., and Kaufman, R.J. (2007). Endoplasmic Reticulum Stress and Oxidative Stress: A Vicious Cycle or a Double-Edged Sword? Antioxid. Redox Signal. 9, 2277–2294.

Marciniak, S.J., Yun, C.Y., Oyadomari, S., Novoa, I., Zhang, Y., Jungreis, R., Nagata, K., Harding, H.P., and Ron, D. (2004). CHOP induces death by promoting protein synthesis and oxidation in the stressed endoplasmic reticulum. Genes Dev. 18, 3066–3077.

Mattaini, K.R., Sullivan, M.R., and Vander Heiden, M.G. (2016). The importance of serine metabolism in cancer. J. Cell Biol. 214, 249–257.

Mckenzie, M., and Ryan, M.T. (2010). Assembly factors of human mitochondrial complex I and their defects in disease. IUBMB Life 62, 497–502.

McManus, J., Cheng, Z., and Vogel, C. (2015). Next-generation analysis of gene expression regulation – comparing the roles of synthesis and degradation. Mol. Biosyst. 11, 2680–2689.

Mendes, C.S., Levet, C., Chatelain, G., Dourlen, P., Fouillet, A., Dichtel-Danjoy, M.-L., Gambis, A., Ryoo, H.D., Steller, H., and Mollereau, B. (2009). ER stress protects from retinal degeneration. EMBO J. 28, 1296–1307.

Mick, D.U., Dennerlein, S., Wiese, H., Reinhold, R., Pacheu-Grau, D., Lorenzi, I., Sasarman, F., Weraarpachai, W., Shoubridge, E.A., Warscheid, B., et al. (2012). MITRAC links mitochondrial protein translocation to respiratory-chain assembly and translational regulation. Cell 151, 1528–1541.

Mignone, F., Gissi, C., Liuni, S., and Pesole, G. (2002). Untranslated regions of mRNAs. Genome Biol. 3, REVIEWS0004.

Morris, A.R., Mukherjee, N., and Keene, J.D. (2008). Ribonomic analysis of human Pum1 reveals cis-trans conservation across species despite evolution of diverse mRNA target sets. Mol. Cell. Biol. 28, 4093–4103.

Mungrue, I.N., Pagnon, J., Kohannim, O., Gargalovic, P.S., and Lusis, A.J. (2009). CHAC1/MGC4504 is a novel proapoptotic component of the unfolded protein response, downstream of the ATF4-ATF3-CHOP cascade. J. Immunol. 182, 466–476.

Munschauer, M. (2015). High-Resolution Profiling of Protein-RNA Interactions (Springer).

Mustoe, A.M., Busan, S., Rice, G.M., Hajdin, C.E., Peterson, B.K., Ruda, V.M., Kubica, N., Nutiu, R., Baryza, J.L., and Weeks, K.M. (2018). Pervasive Regulatory Functions of mRNA Structure Revealed by High-Resolution SHAPE Probing. Cell.

Nakamura, J., Purvis, E.R., and Swenberg, J.A. (2003). Micromolar concentrations of hydrogen peroxide induce oxidative DNA lesions more efficiently than millimolar concentrations in mammalian cells. Nucleic Acids Res. 31, 1790–1795.

Namkoong, S., Ho, A., Woo, Y.M., Kwak, H., and Lee, J.H. (2018). Systematic Characterization of Stress-Induced RNA Granulation. Mol. Cell 70, 175–187.e8.

Oommen, D., and Prise, K.M. (2013). Down-regulation of PERK enhances resistance to ionizing radiation. Biochem. Biophys. Res. Commun. 441, 31–35.

Pakos-Zebrucka, K., Koryga, I., Mnich, K., Ljujic, M., Samali, A., and Gorman, A.M. (2016). The integrated stress response. EMBO Rep. 17, 1374–1395.

Palam, L.R., Baird, T.D., and Wek, R.C. (2011). Phosphorylation of eIF2 facilitates ribosomal bypass of an inhibitory upstream ORF to enhance CHOP translation. J. Biol. Chem. 286, 10939–10949.

Parker, B.J., Moltke, I., Roth, A., Washietl, S., Wen, J., Kellis, M., Breaker, R., and Pedersen, J.S. (2011). New families of human regulatory RNA structures identified by comparative analysis of vertebrate genomes. Genome Res. 21, 1929–1943.

Pascal, R., and Boiteau, L. (2011). Energy flows, metabolism and translation. Philos. Trans. R. Soc. Lond. B Biol. Sci. 366, 2949–2958.

Planet, E., Attolini, C.S.-O., Reina, O., Flores, O., and Rossell, D. (2011). htSeqTools: high-throughput sequencing quality control, processing and visualization in R. Bioinformatics 28, 589–590.

Plemper, R.K., and Wolf, D.H. (1999). Retrograde protein translocation: ERADication of secretory proteins in health and disease. Trends Biochem. Sci. 24, 266–270.

Ponsero, A.J., Igbaria, A., Darch, M.A., Miled, S., Outten, C.E., Winther, J.R., Palais, G., DΆutréaux, B., Delaunay-Moisan, A., and Toledano, M.B. (2017). Endoplasmic Reticulum Transport of Glutathione by Sec61 Is Regulated by Ero1 and Bip. Mol. Cell 67, 962–973.e5.

Pyhtila, B., Zheng, T., Lager, P.J., Keene, J.D., Reedy, M.C., and Nicchitta, C.V. (2008). Signal sequence– and translation-independent mRNA localization to the endoplasmic reticulum. RNA 14, 445–453.

Qiu, C., Kershner, A., Wang, Y., Holley, C.P., WIinski, D., Keles, S., Kimble, J., Wickens, M., and Tanaka Hall, T.M. (2011). Divergence of Pumilio/fem-3mRNA Binding Factor (PUF) Protein Specificity through Variations in an RNA-binding Pocket. J. Biol. Chem. 287, 6949–6957.

Quirós, P.M., Prado, M.A., Zamboni, N., DΆmico, D., Williams, R.W., Finley, D., Gygi, S.P., and Auwerx, J. (2017). Multi-omics analysis identifies ATF4 as a key regulator of the mitochondrial stress response in mammals. J. Cell Biol. 216, 2027–2045.

Rainbolt, T.K., Atanassova, N., Genereux, J.C., and Wiseman, R.L. (2013). Stress-regulated translational attenuation adapts mitochondrial protein import through Tim17A degradation. Cell Metab. 18, 908–919.

Reid, D.W., Chen, Q., Tay, A.S.-L., Shenolikar, S., and Nicchitta, C.V. (2014). The unfolded protein response triggers selective mRNA release from the endoplasmic reticulum. Cell 158, 1362–1374.

Ron, D., and Walter, P. (2007). Signal integration in the endoplasmic reticulum unfolded protein response. Nat. Rev. Mol. Cell Biol. 8, 519–529.

Sammeth, M., Foissac, S., and Guigó, R. (2008). A general definition and nomenclature for alternative splicing events. PLoS Comput. Biol. 4, e1000147.

Scheper, W., and Hoozemans, J.J.M. (2015). The unfolded protein response in neurodegenerative diseases: a neuropathological perspective. Acta Neuropathol. 130, 315–331.

Schönthal, A.H. (2012). Endoplasmic reticulum stress: its role in disease and novel prospects for therapy. Scientifica 2012, 857516.

Schwanhäusser, B., Busse, D., Li, N., Dittmar, G., Schuchhardt, J., Wolf, J., Chen, W., and Selbach, M. (2011). Global quantification of mammalian gene expression control. Nature 473, 337–342.

Sinvani, H., Haimov, O., Svitkin, Y., Sonenberg, N., Tamarkin-Ben-Harush, A., Viollet, B., and Dikstein, R. (2015). Translational tolerance of mitochondrial genes to metabolic energy stress involves TISU and elF1-elF4GI cooperation in start codon selection. Cell Metab. 21, 479–492.

Tchourine, K., Poultney, C.S., Wang, L., Silva, G.M., Manohar, S., Mueller, C.L., Bonneau, R., and Vogel, C. (2014). One third of dynamic protein expression profiles can be predicted by a simple rate equation. Mol. Biosyst. 10, 2850–2862.

Teo, G., Vogel, C., Ghosh, D., Kim, S., and Choi, H. (2014). PECA: A Novel Statistical Tool for Deconvoluting Time-Dependent Gene Expression Regulation. J. Proteome Res. 13, 29–37.

Teo, G., Bin Zhang, Y., Vogel, C., and Choi, H. (2018). PECAplus: statistical analysis of time-dependent regulatory changes in dynamic single-omics and dual-omics experiments. NPJ Syst Biol Appl 4, 3.

Tiwari, S., Askari, J.A., Humphries, M.J., and Bulleid, N.J. (2011). Divalent cations regulate the folding and activation status of integrins during their intracellular trafficking. J. Cell Sci. 124, 1672–1680.

Tolstrup, A.B., Bejder, A., Fleckner, J., and Justesen, J. (1995). Transcriptional regulation of the interferon-gamma-inducible tryptophanyl-tRNA synthetase includes alternative splicing. J. Biol. Chem. 270, 397–403.

Trapnell, C., Roberts, A., Goff, L., Pertea, G., Kim, D., Kelley, D.R., Pimentel, H., Salzberg, S.L., Rinn, J.L., and Pachter, L. (2012). Differential gene and transcript expression analysis of RNA-seq experiments with TopHat and Cufflinks. Nat. Protoc. 7, 562–578.

Tyanova, S., and Cox, J. (2018). Perseus: A Bioinformatics Platform for Integrative Analysis of Proteomics Data in Cancer Research. Methods Mol. Biol. 1711, 133–148.

Tyanova, S., Temu, T., and Cox, J. (2016). The MaxQuant computational platform for mass spectrometry-based shotgun proteomics. Nat. Protoc. 11, 2301–2319.

Uemura, A., Oku, M., Mori, K., and Yoshida, H. (2009). Unconventional splicing of XBP1 mRNA occurs in the cytoplasm during the mammalian unfolded protein response. J. Cell Sci. 122, 2877–2886.

Urra, H., Dufey, E., Lisbona, F., Rojas-Rivera, D., and Hetz, C. (2013). When ER stress reaches a dead end. Biochim. Biophys. Acta 1833, 3507–3517.

Vacaru, A.M., Di Narzo, A.F., Howarth, D.L., Tsedensodnom, O., Imrie, D., Cinaroglu, A., Amin, S., Hao, K., and Sadler, K.C. (2014). Molecularly defined unfolded protein response subclasses have distinct correlations with fatty liver disease in zebrafish. Dis. Model. Mech. 7, 823–835.

Vattem, K.M., and Wek, R.C. (2004). Reinitiation involving upstream ORFs regulates ATF4 mRNA translation in mammalian cells. Proc. Natl. Acad. Sci. U. S. A. 101, 11269–11274.

Vazquez, A., Markert, E.K., and Oltvai, Z.N. (2011). Serine biosynthesis with one carbon catabolism and the glycine cleavage system represents a novel pathway for ATP generation. PLoS One 6, e25881.

Ventoso, I., Kochetov, A., Montaner, D., Dopazo, J., and Santoyo, J. (2012). Extensive translatome remodeling during ER stress response in mammalian cells. PLoS One 7, e35915.

Vervliet, T., Kiviluoto, S., and Bultynck, G. (2012). ER Stress and UPR Through Dysregulated ER Ca2 Homeostasis and Signaling. In Endoplasmic Reticulum Stress in Health and Disease, pp. 107–142.

Vizcaíno, J.A., Csordas, A., del-Toro, N., Dianes, J.A., Griss, J., Lavidas, I., Mayer, G., Perez-Riverol, Y., Reisinger, F., Ternent, T., et al. (2016). 2016 update of the PRIDE database and its related tools. Nucleic Acids Res. 44, D447–D456.

Vogel, C., and Marcotte, E.M. (2012). Insights into the regulation of protein abundance from proteomic and transcriptomic analyses. Nat. Rev. Genet. 13, 227–232.

Vogel, R.O., Janssen, R.J.R.J., Ugalde, C., Grovenstein, M., Huijbens, R.J., Visch, H.-J., van den Heuvel, L.P., Willems, P.H., Zeviani, M., Smeitink, J.A.M., et al. (2005). Human mitochondrial complex I assembly is mediated by NDUFAF1. FEBS J. 272, 5317–5326.

Wakasugi, K., Slike, B.M., Hood, J., Otani, A., Ewalt, K.L., Friedlander, M., Cheresh, D.A., and Schimmel, P. (2002). A human aminoacyl-tRNA synthetase as a regulator of angiogenesis. Proc. Natl. Acad. Sci. U. S. A. 99, 173–177.

Weidmann, C.A., Raynard, N.A., Blewett, N.H., Van Etten, J., and Goldstrohm, A.C. (2014). The RNA binding domain of Pumilio antagonizes poly-adenosine binding protein and accelerates deadenylation. RNA 20, 1298–1319.

Wek, R.C., and Cavener, D.R. (2007). Translational control and the unfolded protein response. Antioxid. Redox Signal. 9, 2357–2371.

Weraarpachai, W., Antonicka, H., Sasarman, F., Seeger, J., Schrank, B., Kolesar, J.E., Lochmüller, H., Chevrette, M., Kaufman, B.A., Horvath, R., et al. (2009). Mutation in TACO1, encoding a translational activator of COX I, results in cytochrome c oxidase deficiency and late-onset Leigh syndrome. Nat. Genet. 41, 833–837.

Wu, X., and Bartel, D.P. (2017). Widespread Influence of 3’-End Structures on Mammalian mRNA Processing and Stability. Cell 169, 905–917.e11.

Xue, S., and Barna, M. (2015). Cis-regulatory RNA elements that regulate specialized ribosome activity. RNA Biol. 12, 1083–1087.

Xue, S., Tian, S., Fujii, K., Kladwang, W., Das, R., and Barna, M. (2014). RNA regulons in Hox 5’ UTRs confer ribosome specificity to gene regulation. Nature 517, 33–38.

Yamamori, T., Meike, S., Nagane, M., Yasui, H., and Inanami, O. (2013). ER stress suppresses DNA double-strand break repair and sensitizes tumor cells to ionizing radiation by stimulating proteasomal degradation of Rad51. FEBS Lett. 587, 3348–3353.

Yang, M., and Vousden, K.H. (2016). Serine and one-carbon metabolism in cancer. Nat. Rev. Cancer 16, 650–662.

Yoshida, H. (2007). ER stress and diseases. FEBS J. 274, 630–658.

Zhao, C., Datta, S., Mandal, P., Xu, S., and Hamilton, T. (2010). Stress-sensitive regulation of IFRD1 mRNA decay is mediated by an upstream open reading frame. J. Biol. Chem. 285, 8552–8562.

Zhao, E., Ding, J., Xia, Y., Liu, M., Ye, B., Choi, J.-H., Yan, C., Dong, Z., Huang, S., Zha, Y., et al. (2016). KDM4C and ATF4 Cooperate in Transcriptional Control of Amino Acid Metabolism. Cell Rep. 14, 506–519.

Zhou, D., Palam, L.R., Jiang, L., Narasimhan, J., Staschke, K.A., and Wek, R.C. (2008). Phosphorylation of eIF2 directs ATF5 translational control in response to diverse stress conditions. J. Biol. Chem. 283, 7064–7073.

Zhou, X., He, L., Wu, C., Zhang, Y., Wu, X., and Yin, Y. (2017). Serine alleviates oxidative stress via supporting glutathione synthesis and methionine cycle in mice. Mol. Nutr. Food Res. 61.

Zito, E., Melo, E.P., Yang, Y., Wahlander, Å., Neubert, T.A., and Ron, D. (2010). Oxidative protein folding by an endoplasmic reticulum-localized peroxiredoxin. Mol. Cell 40, 787–797.

